# The Septin Cytoskeleton is a Novel Regulator of Intestinal Epithelial Barrier Integrity and Mucosal Inflammation

**DOI:** 10.1101/2024.12.20.629767

**Authors:** Nayden G. Naydenov, Gaizun Hu, Atif Zafar, Dominik Robak, Khosiyat Makhmudova, Susana Lechuga, Yuta Ohno, Naseer Sagwan, Saikat Bandyopadhyay, Ryan Musich, Erin Jeffery, Lei Sun, Armando Marino-Melendez, Florian Rieder, Gloria Sheynkman, Andrei I. Ivanov, Seham Ebrahim

**Affiliations:** Department of Inflammation and Immunity, Lerner Research Institute, Cleveland Clinic Foundation, Cleveland, OH 44195; Department of Molecular Physiology and Biological Physics, School of Medicine, University of Virginia VA; Department of Cardiovascular and Metabolic Science, Lerner Research Institute, Cleveland Clinic Foundation, Cleveland, OH 44195; Department of Gastroenterology, Hepatology and Nutrition, Digestive Diseases Institute, Cleveland Clinic Foundation, Cleveland, OH 44195

**Keywords:** SEPT9, epithelial barrier, cytoskeleton, colitis, mucosal inflammation, myosin II, tight junctions

## Abstract

**Background and Aims:** Intestinal epithelial barrier-integrity is essential for human health, and its disruption induces and exacerbates intestinal inflammatory disorders. While the cytoskeleton is critical for maintaining gut barrier-integrity, the role of the septins- the newest family of cytoskeletal proteins- is unknown. To address this knowledge gap, we evaluate the role of SEPT9- a critical component of the septin-cytoskeleton- in intestinal epithelial cell (IEC) barrier permeability and inflammation.

**Methods:** We developed SEPT9-NeonGreen knockin mice, inducible intestinal epithelial cell (IEC)-specific SEPT9 knockout (KO) mice, and SEPT9-KO human IEC lines. SEPT9 localization was analyzed using super-resolution microscopy. Barrier-integrity was assessed via transepithelial electrical resistance, FITC-dextran flux, and visualization of tight junction (TJ) and adherens junction (AJ) proteins. Dextran sodium sulfate-induced experimental colitis was evaluated in control and KO mice through measuring cytokine expression, immune cell infiltration, and IEC death. SEPT9 expression was examined in intestinal tissue of IBD patients.

**Results:** SEPT9 overlapped with TJs and AJs at IEC apical junctions. SEPT9 loss resulted in a leaky epithelial barrier due to mislocalization of junctional proteins. SEPT9 interacted with non-muscle myosin IIC (NMIIC) at the IEC apical-junctional actomyosin belt, and its ablation displaced NMIIC from IEC junctions. Loss of NMIIC also caused barrier disruption. SEPT9 KO mice exhibited increased susceptibility to experimental-colitis. SEPT9 expression was significantly reduced in intestinal mucosa of IBD patients.

**Conclusion:** SEPT9 regulates intestinal barrier integrity, supporting TJ and AJ assembly through NMIIC recruitment to the actomyosin belt. SEPT9 safeguards the intestinal mucosa during acute inflammation, and its reduced expression in IBD suggests a loss of this protective function.

**Graphical Abstract:** 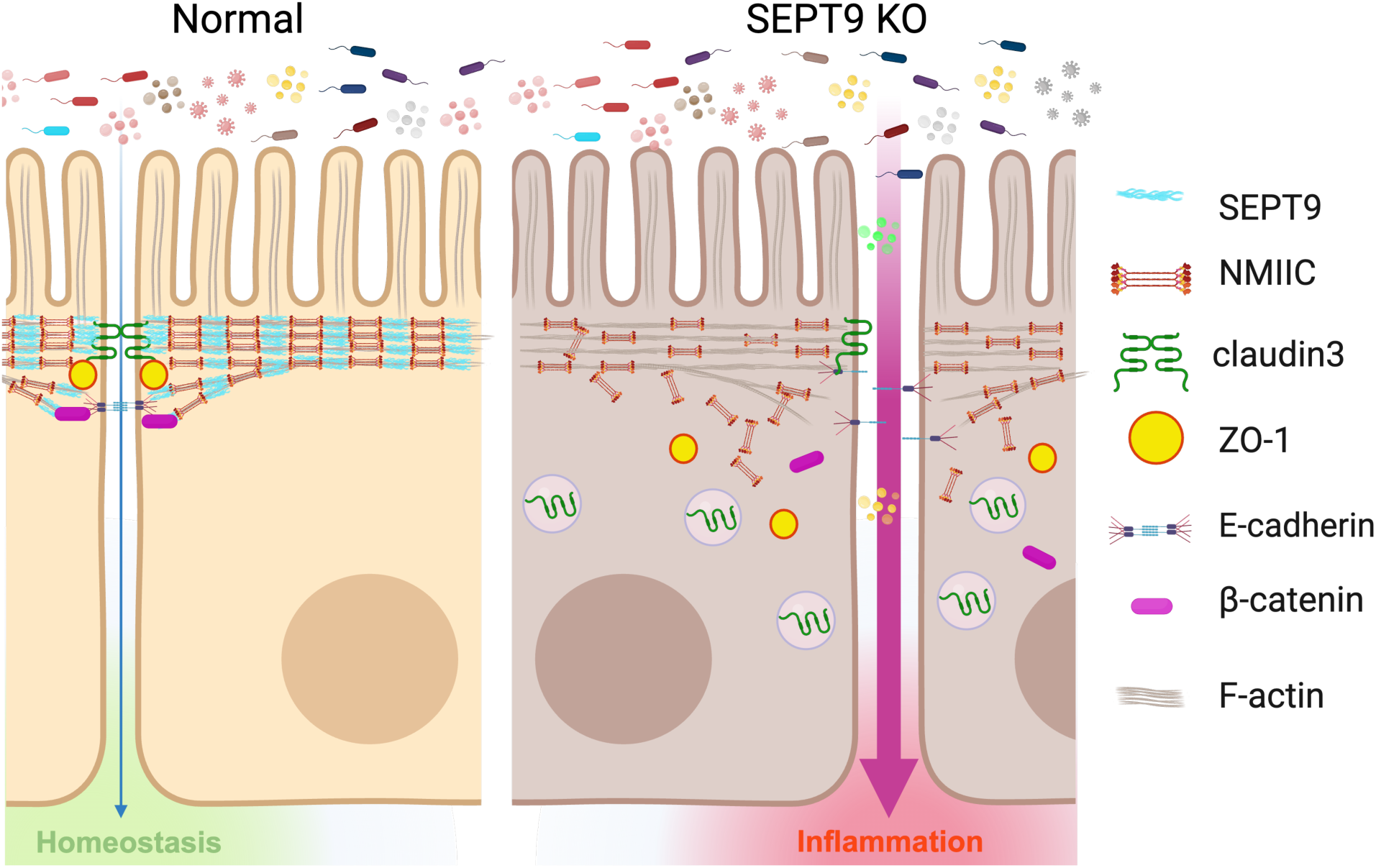

## INTRODUCTION

The intestinal epithelial barrier is a critical component of gut health, forming a protective boundary that separates internal organs from luminal contents, including microorganisms, while regulating the directional flow of water, nutrients, and waste. Disruption of this barrier is a hallmark of various gastrointestinal disorders, most notably inflammatory bowel diseases (IBD) such as Crohn’s disease (CD) and ulcerative colitis (UC)^1–3^. These barrier defects are key drivers of disease progression, triggering and sustaining mucosal inflammation^3–5^. Gut barrier is composed of a single layer of columnar epithelial cells interconnected by lateral cell-cell adhesions, mediated by specialized multiprotein complexes known as epithelial junctions. The three primary types of epithelial junctions are tight junctions (TJs), adherens junctions (AJs), and desmosomes. Among these, the apically located TJs play a central role in maintaining epithelial barrier integrity and regulating permeability, underscoring their significance in gut health and disease^6–9^.

TJs are formed via self-assembly of various transmembrane proteins, including claudins, occludin, and junctional adhesion molecule A^6–8^. These proteins interact with counterparts on adjacent plasma membranes, establishing a selective barrier. On their cytoplasmic side, TJs are anchored by members of the zonula occludens (ZO) protein family, along with scaffolding, polarity, and signaling proteins^6–8^. TJs work in tandem with E-cadherin-based adherens junctions (AJs) to regulate epithelial barrier integrity and facilitate remodeling^9,10^. A defining feature of TJs and AJs is their coupling to the circumferential actomyosin belt, composed of antiparallel actin filament bundles and non-muscle myosin II (NMII) motors^11–13^. This connection to the actomyosin cytoskeleton is essential for maintaining junction stability, regulating permeability, and enabling dynamic remodeling^11–13^.

Interestingly, super-resolution microscopy has revealed that the peri-junctional NMII assembles into periodic, sarcomere-like structures^14,15^, suggesting a critical role in mechanical force transduction and the regulation of junctional permeability^16^. Despite these insights, the mechanisms governing the recruitment and assembly of the junction-associated actomyosin cytoskeleton, particularly in healthy versus inflamed intestinal epithelium, remain poorly understood. This gap in knowledge underscores the need to further explore how cytoskeletal dynamics contribute to epithelial barrier function in both intestinal homeostasis and disease.

Septin GTPases are a family of filament-forming proteins with the distinctive ability to interact with cellular membranes and major cytoskeletal components, such as actin filaments and microtubules^17–19^. Septins have been reported to cross-link and curve actin filaments^20,21^, anchor actin bundles to the plasma membrane^22^, interact with NMII and regulate its activity^21,23^. These properties make the septin cytoskeleton a compelling candidate for controlling the assembly and contractility of the peri-junctional actomyosin belt, thereby playing a potential role in maintaining epithelial barrier integrity. While some septins were previously implicated in the regulation of epithelial and endothelial junctions *in vitro*^24–26^, the underlying molecular mechanisms and *in vivo* relevance of the septin-dependent control of tissue barrier integrity remain enigmatic.

This represents an important knowledge gap, especially given the reported association between septin dysregulation, specifically septin 9 (SEPT9), and human intestinal disorders. Both mutations and hypermethylation of the SEPT9 gene, resulting in reduced protein expression, occur in colorectal cancer (CRC)^27,28^. Since colon carcinogenesis is known to be modulated by gut inflammation and inflammation-induced disruption of the intestinal epithelial barrier^29,30^, this may have relevance on how SEPT9-based cytoskeleton regulates the intestinal epithelial barrier integrity in the setting of mucosal inflammatory disorders.

In this study, we investigate the cellular localization and function of SEPT9 in the intestinal epithelium using a combination of models, including knockin mice expressing fluorescently-tagged endogenous SEPT9, epithelial-specific SEPT9 knockout mice, and SEPT9-deficient human intestinal epithelial cells (IEC). We demonstrate, for the first time, that SEPT9 is enriched at epithelial apical junctions, colocalizing with TJ and AJ proteins. SEPT9 regulates intestinal epithelial barrier integrity via the recruitment of junctional NMIIC and assembly of TJ components. Furthermore, SEPT9 plays protective roles during experimental colitis *in vivo* by limiting mucosal injury and inflammation. Consistent with these findings, epithelial SEPT9 expression is diminished in the intestinal mucosa of IBD patients and is mislocalized in IEC of Crohn’s disease patients. Our findings thus reveal a novel role for septin cytoskeleton in regulating epithelial junctional integrity *in vivo*, unravel the mechanism underlying the barrier-protective function of SEPT9, and highlight that abnormal organization of the septin cytoskeleton in IBD patients could be an important contributor to chronic mucosal inflammation.

## METHODS

### Animals

Mice from C57BL/6J backgrounds were kept in the animal care facilities at the University of Virginia or Lerner Research Institute under controlled temperature and humidity using 12:12-hour light-to-dark cycles. Animals were supplied with water and food *ad libitum*. All involved animals were reviewed and received approval from the Animal Welfare Committee of the University of Virginia and Lerner Research Institute. See Supplementary Methods for the details of tamoxifen-inducible, intestinal epithelium-specific knockout mice (SEPT9-KO) and SEPT9-mNeonGreen reporter mice.

### Human IBD samples

Deidentified full thickness intestinal surgical resection specimens obtained from inflammatory bowel diseases (IBD) patients with active Crohn’s Disease (CD), ulcerative colitis (UC) and non-IBD controls (diverticulosis or non-affected margins of resected colorectal cancer) were provided by the Human Tissue Procurement Facility of the Cleveland Clinic through the services of the Biorepository Core. The human tissue collection was approved by Cleveland Clinic Institutional Review Board (IRB) under minimal risk IRB approval number 17-1167. The human tissue harvested at the University of Virginia was approved by the University of Virginia’s Institutional Biosafety Committee (IBC) (protocol number: 9987-22). Patients provided informed consent for the tissue sample collections. The demographic information for different patient groups is provided in Supplementary Table 1.

### Dextran sodium sulfate (DSS) colitis

Experimental colitis was induced in 8–10 week-old SEPT9-KO mice and control littermates by administering a 3% (w/v) solution of DSS in drinking water. Unchallenged animals received tap water *ad libitum*. Animals were weighed daily and monitored for signs of intestinal inflammation. The disease activity index was calculated by averaging numerical scores of body weight loss, stool consistency, and intestinal bleeding. Detailed information about diseases activity index, sample collection and processing is provided in the Supplementary Methods. The *in vivo* intestinal permeability assay was performed in SEPT9-KO and control mice under normal conditions and after DSS colitis by administering different size molecular tracers. For details see Supplementary methods.

### Generation of SEPT9 and NMIIC knockout cell lines

SEPT9 knockout HT-29 and NMIIC-knockout Caco-2BBE cell lines were constructed using the CRISPR/Cas9 V2 system. Single guide RNAs (sgRNA) were designed with a CRISPR design website (http://crispr.mit.edu/), provided by the Feng Zhang Lab (McGovern Institute, Massachusetts Institute of Technology, Boston, MA, USA). Their sequences are as follows: SEPT9 sg4-ACGGAACGAGAAGGCCCCGG, sg6-AGAGGGTCCACTTCAAACAG, NMIIC sg1-CCTCGGTCACTGCGTACACG and control sg-GACCGGAACGATCTCGCGTA. SEPT9, NMIIC and control sgRNAs were cloned into a BbsI restriction site of the lentiCRISPR v2 vector (Addgene Watertown, MA, Cat# 52961) and confirmed by sequencing. Lentiviruses were produced in HEK293T cells co-transfected with packaging plasmids pCD/NL-BH*DDD (Addgene, Cat# 17531,) and pLTR-G (Addgene, Cat# 17532) using a TransIT-293 transfection reagent (Mirus Bio, Madison, WI, USA). Stable SEPT9 and NMIIC-depleted IEC cells were generated by transducing HT-29 and Caco-2BBE cells respectively with lentiviruses containing corresponding sgRNAs and subsequent puromycin selection (5 μg/mL) for 7 days. Control IEC were generated via transduction with the control sgRNA containing lentivirus and puromycin selection.

### Transepithelial electrical resistance (TEER) and FITC dextran permeability measurements in vitro

IECs were seeded on the 24-well Transwell® membrane inserts with 3-micron pore size (Corning, Cat# 3415) coated with rat tail collagen I. The TEER was measured using an epithelial ohmmeter (EVOM2, World Precision Instruments, Sarasota, FL). The resistance of cell-free collagen-coated filters was subtracted from each experimental point. *In vitro* FITC-dextran permeability assay was performed at indicated times as previously described^9^. IEC monolayers cultured on Transwell filters were apically exposed to 1 mg/ml of FITC-labeled dextran (4 kDa) in HEPES-buffered Hanks’ balanced salt solution (HBSS+). After 120 min of incubation, samples were collected from the lower chamber, and FITC fluorescence intensity was measured using a BioTek Synergy H1 plate reader, at excitation and emission wavelengths 485 nm and 544 nm, respectively. The amount of FITC-dextran translocated across the epithelial cell monolayer was calculated based on a calibration curve plotted with serial dilutions of the FITC-dextran stock.

### Immunofluorescence labeling and microscopy of whole mouse colon

For whole tissue staining for *en face* imaging, fixed whole tissues were rinsed and cut to the desired size. The tissue samples were then permeabilized in 0.5% Triton X-100-PBS solution (PBST) for 45∼60min at room temperature. The tissues were blocked in 10% normal goat serum for 2 hours at room temperature and then incubated with primary antibodies overnight at 4°C. Next, the tissues were labeled with AlexaFluor-conjugated secondary antibodies for 1 hour at room temperature. Confocal microscopy was conducted using a CSU-W1 Yokogawa spinning disk field scanning confocal system. For image acquisition from fixed tissue sections, a sampling frequency of 30–60 nm was achieved at the camera sensor. Image acquisition was controlled with NIS-Elements software (Nikon Instruments). For bright-field microscopy, SEPT9 immunolabeled samples were imaged using a THUNDER imager system (THUNDER X Imager, Leica) with large image alignment.

### Image processing and segmentation

Machine learning-based morphological analysis of images stained with junctional markers was conducted using a customized U-Net model implemented in PyTorch^31^. This model was specifically trained and optimized to segment epithelial cells based on their junctional staining patterns. Following the generation of binary segmentation masks, the resulting data were imported into ImageJ (v1.54f)^32^ for quantitative analysis. Individual segmented cells were then inspected, and any cells displaying suboptimal segmentation were manually excluded from further analyses. The remaining high-quality segmentations were used to quantify cell shape descriptors, including cell area, circularity, and solidity, as well as the average fluorescence intensity (FI). To streamline this process, we implemented customized macros that automated the quantification across all segmented images. See Supplementary Methods for detailed description of image analysis for all other immunolabeling and confocal microscopy studies.

### Data processing and statistical analysis

The quantified data from ImageJ was then imported into Python (v3.10) for further processing and statistical analysis. We developed custom Python scripts to gather and process the data efficiently. Subsequently, this processed data was input into GraphPad Prism (v10.4.0) for statistical analysis and visualization. All scripts and code utilized in this study are available upon request. Data are displayed as a mean ± SEM. The statistical significance of the difference between 2 sets of data was evaluated using the two-tailed unpaired Student’s t-test when data were distributed normally.

## RESULTS

### SEPT9 is expressed in different subtypes of intestinal epithelial cells and localizes at epithelial junctions

A publicly available single-cell RNA dataset (GSE242087) was used to examine SEPT9 expression in different cell types in healthy human ileum and colon. SEPT9 was expressed in all major cell populations, including epithelial, immune, and stromal cells (Figure 1A). A deeper examination of different IEC subtypes revealed SEPT9 expression in all major differentiated and progenitor IECs in normal ileal and colonic mucosa being especially abundant in absorptive enterocytes (EC3, EC4), enterochromaffin cells (EEC1, EEC2), transient amplified cells (TA1-3) and unidentified IEC population (Clus12) (Figure 1B). SEPT9 mRNA levels show positive correlation with the expression of other major septins, SEPT2, SEPT7 and SEPT11, in different ileal and colonic IEC subtypes (Supplementary Figure 1A and B), which could indicate co-regulation and possible functional cooperation of these septin paralogs in human intestinal epithelium.

**Fig. 1:**
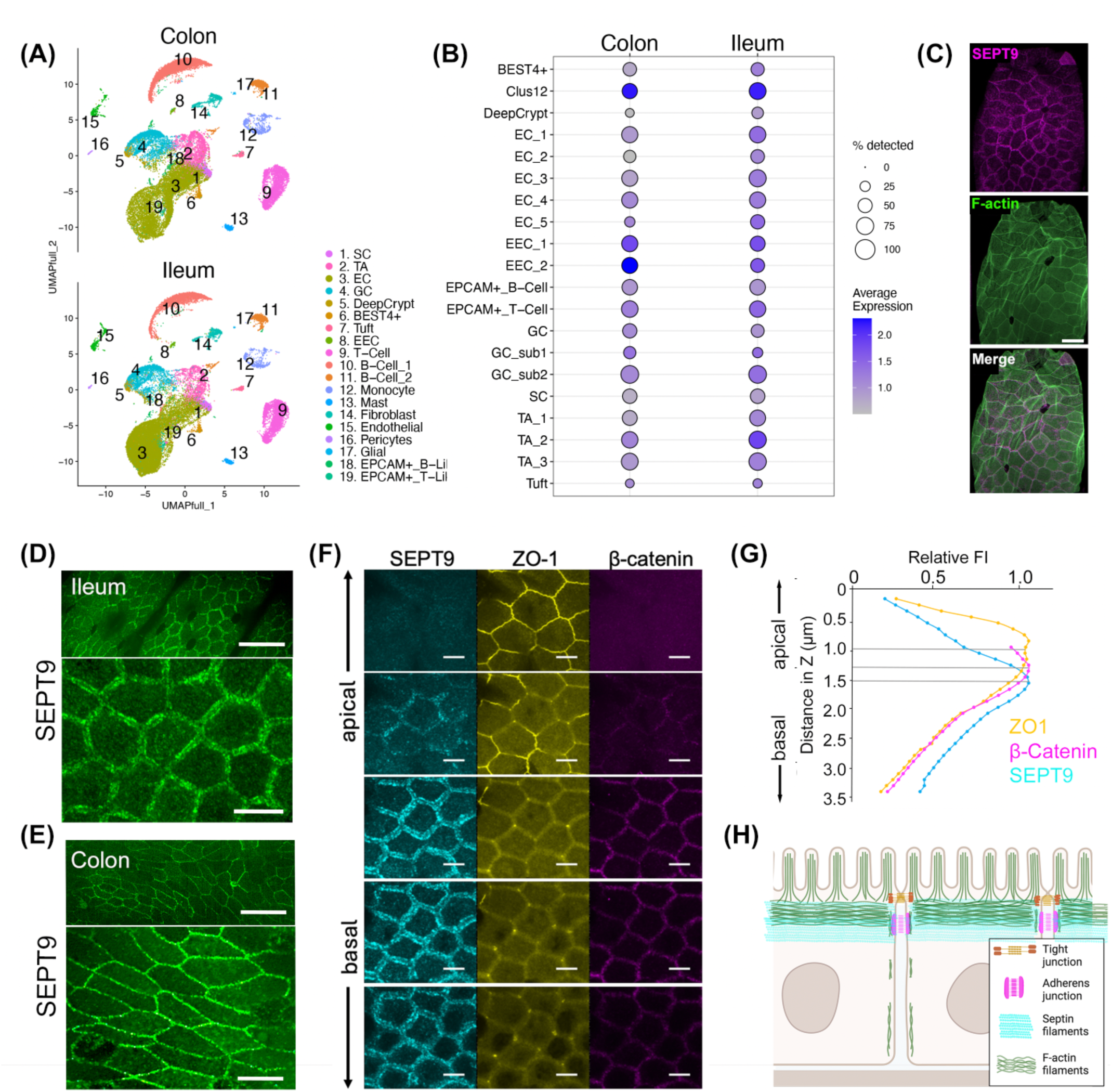
SEPT9 localizes at apical junctions of the intestinal epithelium *in vivo*. Single-cell RNA sequencing analysis of normal ileal and colonic tissues showing **(A)** SEPT9 expression in different types of intestinal cells and **(B)** its distribution in different IEC subtypes. **(C)** Immunofluorescence labeling of whole mount mouse small intestinal tissues with an anti-SEPT9 antibody (magenta) and F-actin probe phalloidin (green). Scale bar= 20 μm. **(D and E)** Localization of endogenous SEPT9 in the ileum and colon of SEPT9-mNG mice. Scale bars= 50 μm in upper and 10 μm in lower images. **(F)** Colocalization of endogenous SEPT9-NeonGreen with immunostained TJ (ZO-1) and AJ (β-catenin) markers in wholemount mouse ileal mucosa. Scale bar= 5 μm. **(G)** Fluorescent intensity plots of SEPT9 (cyan), ZO-1 (yellow) and β-catenin (magenta) at different Z-planes at the IEC apical junctions. **(H)** A schematic representation of SEPT9 localization at apical junctions in IEC.

The subcellular localization of endogenous SEPT9 in mouse intestinal mucosa was evaluated by immunofluorescence labeling in wild-type mice (Figure 1C) as well as in a knockin mice expressing mNeonGreen-tagged SEPT9 (SEPT9^mNG^) (Figure 1D-G). In both cases, confocal microscopy revealed SEPT9 is enriched at epithelial apical junctions in mouse ileum (Figure 1C and D) and colon (Figure 1E). To delineate the specific localization of SEPT9 relative to the IEC junctional complexes, we immunolabeled ileal tissue from SEPT9^mNG^ mice for the major TJ protein, ZO-1, and the AJ protein, β-catenin. We acquired z-stacks of confocal images of the apical junctional region of the ileal epithelium at 200 nm intervals (a subset of these images is shown in Figure 1F). To estimate the relative localization of each tagged protein along the z-axis, we plotted the immunofluorescence intensity values and determined the points of fluorescence maxima. The immunofluorescence signal of ZO-1 along the z-axis was most apical (Figure 1G), while β-catenin and SEPT9 were both more basal. Of note, the fluorescence intensity of SEPT9^mNG^ overlapped partially with both ZO-1 and β-catenin immunofluorescence (Figure 1G). These results are consistent with a SEPT9 network that overlaps with both TJs and AJs (Figure 1H).

### Loss of intestinal epithelial SEPT9 increases gut barrier permeability and alters junctional protein localization

Given the observed association of SEPT9 complexes with mouse intestinal AJs and TJs, we next sought to investigate a potential role for SEPT9 in regulating gut barrier integrity using SEPT9^fl/fl^Vil1^CreERT2^ mice (referred to hereafter as SEPT9-KO), to enable tamoxifen-inducible IEC-specific knockout of SEPT9. Immunoblotting analysis of isolated IECs demonstrated an almost complete loss of SEPT9 protein after tamoxifen-induced *Cre* recombinase activation (Figure 2A and B). This loss of SEPT9 was also validated by *en face* imaging of immunolabeled colonic mucosa of SEPT9-KO animals (Figure 2C).

**Fig. 2:**
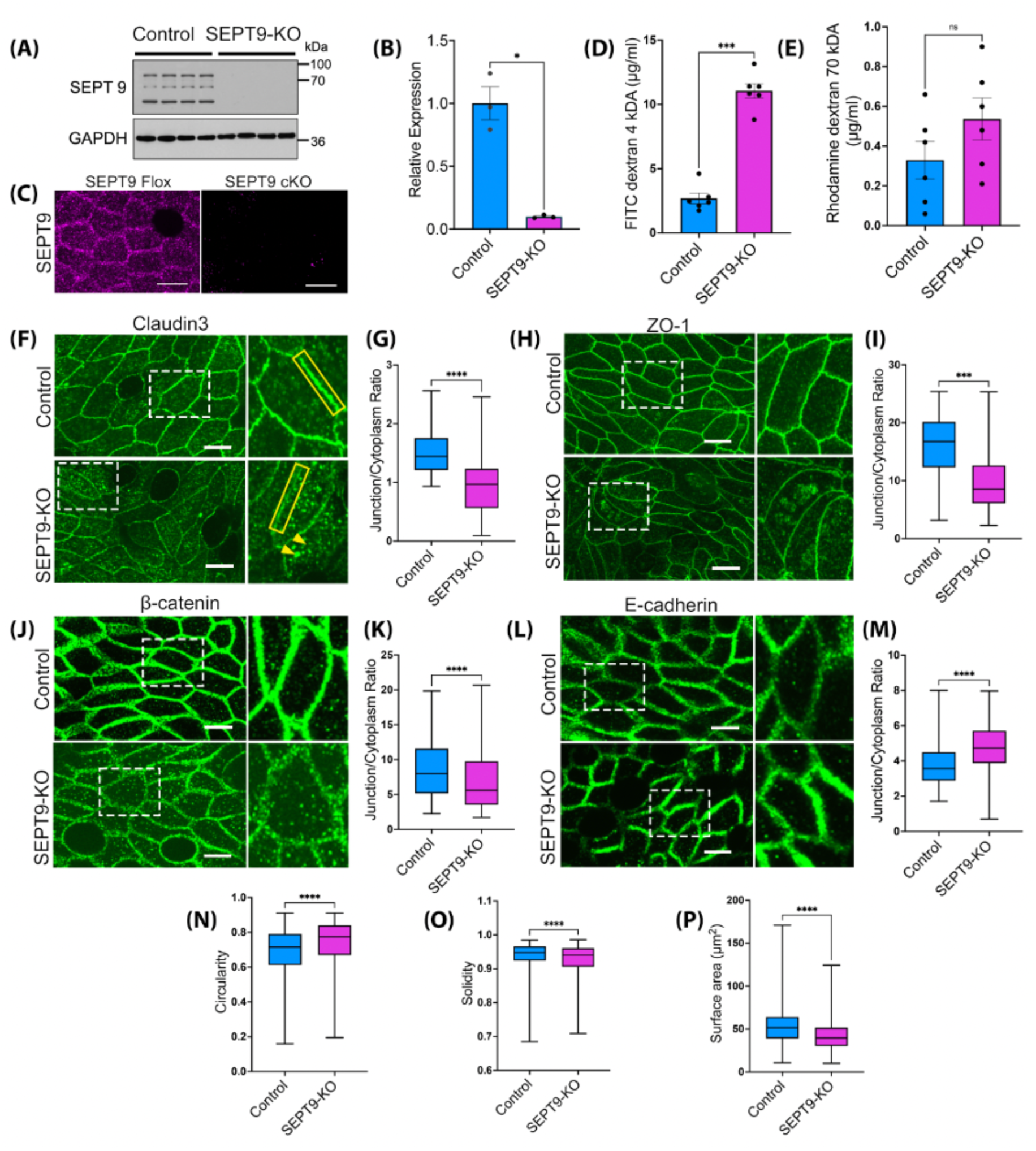
Loss of SEPT9 increases epithelial barrier permeability and impairs apical junctional assembly in the intestinal mucosa *in vivo*. (A-C) Validation of SEPT9 loss in SEPT9-KO mice. Representative immunoblots (A) and densitometric quantification (B) of SEPT9 expression in colonic scrapes, and immunolabeling for SEPT9 in ileal epithelia (C). Mean ± SEM, n= 4; *p<0.05. Scale bar= 10μm. **(D and E)** Gut-to-blood passage of 4 kDa FITC-dextran and 70 kDa Rhodamine-dextran in SEPT9-KO and control mice. Mean ± SEM, n= 6; ****p<0.0001. Representative *en face* images and quantification of immunolabeled junctional markers in wholemount colon of SEPT9-KO and control mice, including claudin3 (F and G), ZO-1 (H and I), β-catenin (J and K) and E-cadherin (L and M). Scale bars= 10 μm. White dash boxes indicate the zoomed areas. Arrowheads highlight accumulation of cytoplasmic TJ and AJ proteins in SEPT9-KO mice. **(N-P)** Measurements of junctional morphology in colonic mucosa of SEPT9-KO and control mice examined via measuring circularity (N), solidity (O), and the cell surface area (P); n= 5/group. Data represented as box-whisker plots with whiskers extending to the minimum and the maximum and mean at the middle of the box body. **p<0.01, ****p<0.0001.

SEPT9-KO mice did not show growth retardation or symptoms of spontaneous gastrointestinal disorders, such as diarrhea, rectal prolapses, or bleeding (data not shown). Consistently, they did not show gross abnormalities of their ileal and colonic mucosa (Supplementary Figure 2A), except for the increase in Goblet cell number (Supplementary Figure 2B-D). Interestingly, a fluorescent tracer-based gut barrier permeability assay showed approximately 4-fold increase in the transmucosal flux of 4 kDa FITC-dextran in SEPT9-KO mice as compared to the control littermates, thereby indicating a leaky gut barrier (Figure 2D). Intestinal permeability to a large tracer, 70 kDa Rhodamine-dextran, was not affected, suggesting that SEPT9 knockout does not lead to nonspecific loss of IECs (Figure 2E).

We hypothesized that the observed increase in intestinal epithelial permeability can be attributed to dysfunctional IEC junctions, particularly TJs in SEPT9-KO mice. Therefore, we used immunofluorescence labeling together with machine-learning image analysis (Figure 2F-M, Supplementary Figure 3A and B) to examine the localization of key TJ and AJ proteins in mouse colonic mucosa. We observed mislocalization of TJ proteins claudin 3 and ZO-1 in IEC of SEPT9-KO mice, with decreased junctional enrichment of these proteins and their increased accumulation in small cytoplasmic vesicles (Figure 2F-I). AJ proteins, β-catenin and E-cadherin were differentially impacted by loss of SEPT9, with β-catenin translocated from IEC junctions into cytoplasmic vesicles (Figure 2J and K) and increased junctional accumulation of E-cadherin (Figure 2 L and M). These observations are consistent with previous studies reporting increased internalization of TJ proteins in animal models with defective intestinal epithelial barrier and mucosal biopsies from IBD patients^33–36^. Of note, the described functional and structural defects of apical junctions in the colonic mucosa of SEPT9-KO mice were not due to altered expression of key AJ and TJ proteins (Supplementary Figure 3C and D).

In addition to mislocalization of junctional proteins *in vivo*, loss of SEPT9 caused noticeable alterations in IEC morphology *in vivo* (Figure 2N-P). The shape of epithelial cell-cell contacts and the epithelial cell size depended on the tension in cell-cell junctions generated by the peri-junctional actomyosin belt^37,38^. We thus posited that alterations in the shape and size of SEPT9 depleted IEC cells could be a consequence of dysregulated junctional tension, which also directly impact epithelial barrier permeability^16^.

To establish whether the observed disruption of the gut barrier represents an autonomous effect of SEPT9 depletion in IEC, we created HT-29 cF8 human colonic epithelial cell lines with CRISPR/Cas9-mediated knockout of SEPT9 (Figure 3A). Similar to the results obtained in murine intestinal mucosa, SEPT9 deficient IEC monolayers demonstrated significantly increased paracellular permeability based on TEER (Figure 3B) and FITC-dextran flux (Figure 3C) measurements. Consistent with the *in vivo* effects of SEPT9 depletion, loss of SEPT9 expression decreased junctional accumulation of claudin 3, ZO-1 and β-catenin (Figure 3D-K), but did not affect expression levels of different AJ/TJ proteins (Supplementary Figure 4A).

**Fig. 3:**
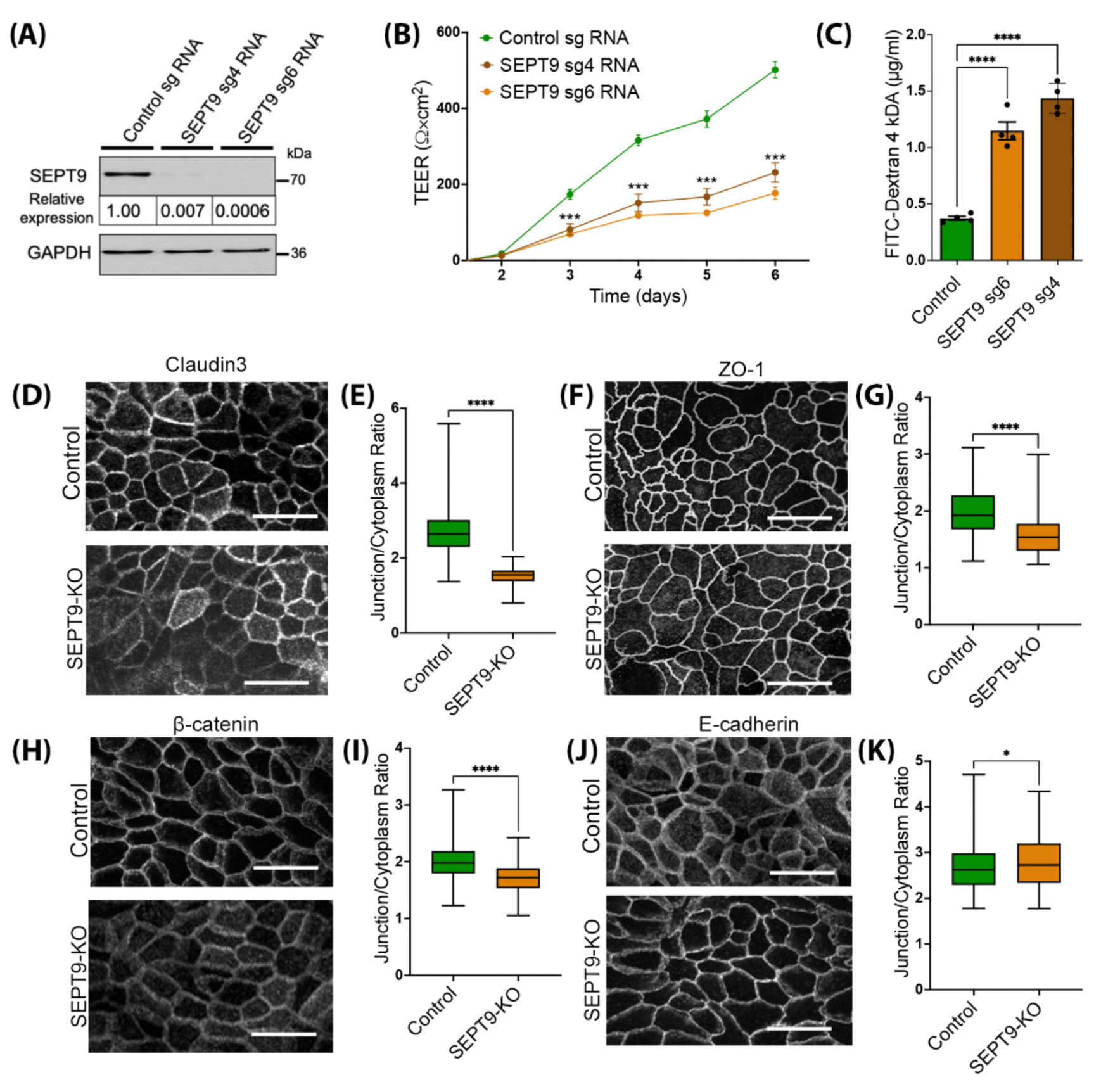
SEPT9 loss increases epithelial barrier permeability and alters the assembly of IEC apical junctions *in vitro*. **(A)** Immunoblotting analysis of SEPT9 expression in HT-29 cells with CRISPR/Cas9 mediated knockout of SEPT9 using two different sgRNAs. **(B)** Transepithelial electrical resistance (TEER) of control and SEPT9-KO HT-29 cells. **(C)** FITC-dextran flux in control and SEPT9-KO HT-29 cells. Mean ± SEM, n= 4; **(D-K)** Representative images and quantification of immunolabeled junctional markers in control and SEPT9-KO HT-29 cells, including claudin3 (D and E), ZO-1 (F and G), β-catenin (H and I), and E-cadherin (J and K). Scale bar= 10 μm. Data represented as box-whisker plots with whiskers extending to the minimum and the maximum and mean at the middle of the box body. **p<0.01, ****p<0.0001.

We next sought to assess the dynamics of TJ proteins as a readout of TJ stability. To do this, we expressed GFP-tagged claudin 7 in control and SEPT9 deficient HT-29 cells and performed live cell imaging. Strikingly, we found that while claudin 7 was primarily localized at apical junctions in control cells, it was translocated into numerous cytoplasmic vesicles after SEPT9 depletion (Supplementary Movie 1). Importantly, SEPT9 knockout in HT-29 cells did not affect IEC proliferation (Supplementary Figure 4B and C), thereby indicating specific regulation of IEC junctions by the SEPT9-based cytoskeleton.

### SEPT9 interacts with non-muscle myosin IIC and regulates junctional localization of this actin motor

To investigate the mechanisms of SEPT9-dependent regulation of the intestinal epithelial barrier permeability and TJ assembly, we next determined the interactome of SEPT9 in IECs by performing Affinity Purification Mass Spectrometry (APMS) on pulldowns of SEPT9-GFP overexpressed in DLD-1 cells. In addition to interactions with several TJ and AJ proteins, analysis of the interactome revealed a strong association of IEC SEPT9 with several major components of the actomyosin cytoskeleton, including non-muscle myosin II motors, NMIIC and NMIIB (Supplementary Figure 5A). These findings, together with our data showing altered IEC shapes in the SEPT9-KO, which could be due to altered actomyosin tension, as well as reported interactions between septins and NMII in different mammalian cells^23,39,40^, led us to hypothesize that loss of SEPT9 causes a reduction in NMII-associated forces at IEC apical junctions, leading to impaired TJ assembly. Since NMIIC is highly enriched^41^, while NMIIB is not expressed^35^, in the intestinal epithelium *in vivo*, we focused on investigating the interactions between SEPT9 and NMIIC.

Super-resolution microscopy revealed a precise and periodic pattern of SEPT9 interdigitating with NMIIC ‘sarcomeres’ at IEC apical junctions (Figure 4A and B). Co-expression of SEPT9 and NMIIC in COS7 cells that lack endogenous NMIIC, resulted in incorporation, and co-localization, of both cytoskeletal proteins into F-actin bundles (Supplementary Figure 5B). As further validation of interaction between SEPT9 and NMIIC *in vivo* and *in vitro*, we observed a significant reduction in junctional NMIIC localization in the intestinal epithelium of SEPT9-KO mice (Figure 4C and D) and SEPT9 deficient HT-29 cell monolayers (Figure 4E and F), which was not accompanied by the altered total cellular levels of NMIIC and NMIIA (Supplementary Figures 3 and 4). These data highlight the selective association of SEPT9 and NMIIC at IEC junctions. Finally, CRISPR-Cas9 dependent knockout of NMIIC in Caco-2 cells (Figure 4G) recapitulated the effects of SEPT9 deletion, resulting in epithelial barrier disruption as measured by significantly decreased TEER (Figure 4H) and increased FITC-dextran flux (Figure 4I). Overall, these data suggest that SEPT9 enhances the integrity of the IEC barrier by recruiting/stabilizing NMIIC motor at epithelial TJs.

**Fig. 4:**
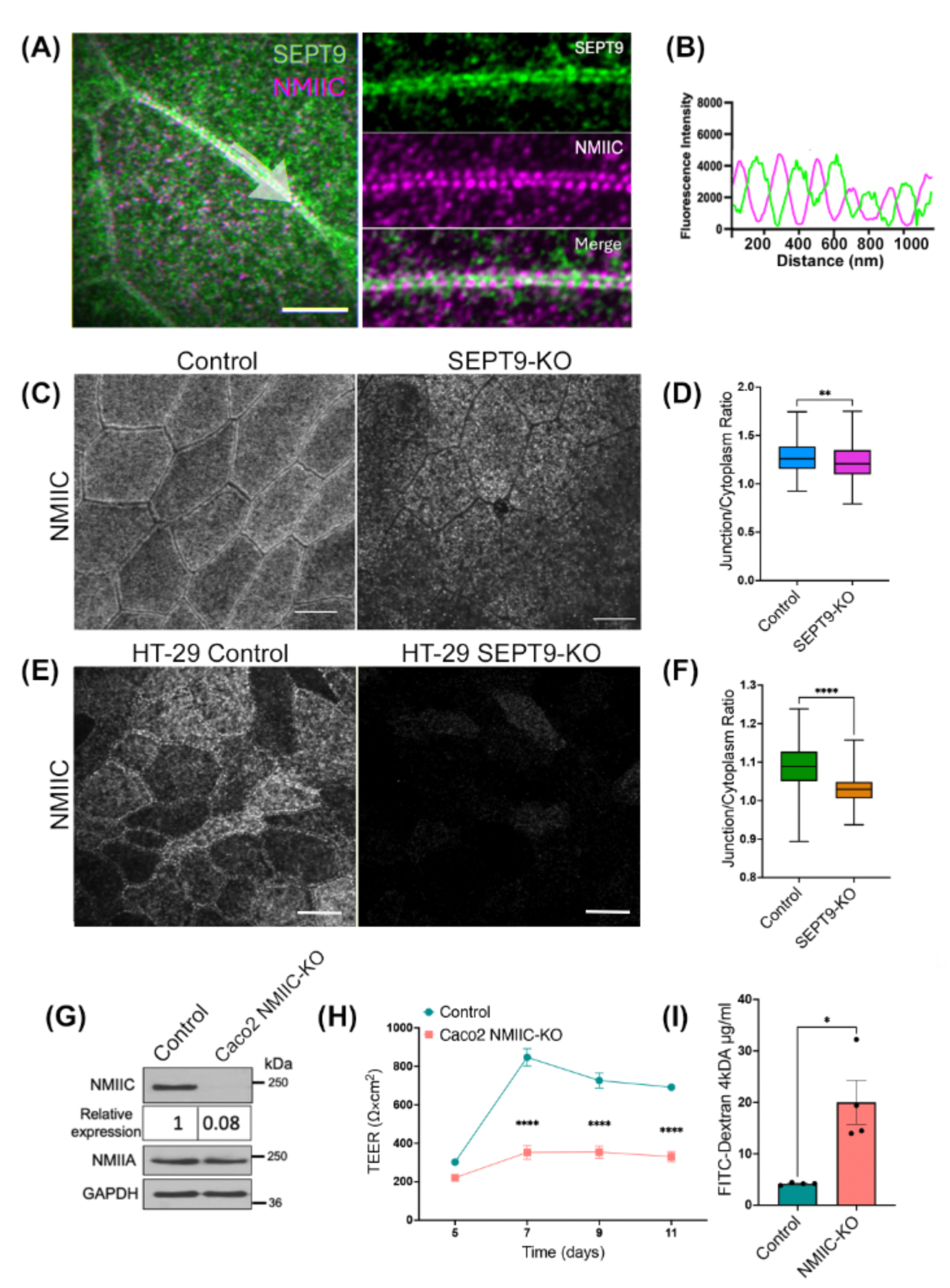
The NMIIC motor is recruited to apical junction by SEPT9 and regulates permeability of the model IEC barrier. **(A)** Super-resolution microscopy image of SEPT9 (green) and NMIIC (magenta) in DLD-1 human colonic epithelial cells. **(B)** Fluorescence intensities profiles show SEPT9 and NMIIC signal intercalation along the cell-cell junction highlighted by the white arrow. **(C and D)** Representative image and junction to cytoplasmic ratio of immunolabeled NMIIC in colonic mucosa of SEPT9-KO and control mice, **(E and F)** Representative image and junction to cytoplasmic ratio of immunolabeled NMIIC in control and SEPT9-KO HT-29 cells. Data presented as box-whisker plots with n= 5/group. **p<0.01, ****p<0.0001; Scale bars= 10 μm. (**G**) Immunoblotting analysis of NMIIC expression in Caco-2 cells with CRISPR/Cas9-mediated knockout of NMIIC. (**H**) TEER of control and NMIIC-KO cells. (**I**) FITC-dextran flux in control and NMIIC-KO Caco-2 cells. Mean ± SEM, n= 4; **p<0.01, **** p<0.001.

### IEC-specific knockout of SEPT9 alters the transcription program in intestinal epithelium in vivo

Given our findings of increased intestinal permeability in unchallenged SEPT9 knockout mice (Figure 2D) we next asked if such barrier leakiness was sufficient to trigger alterations in IEC homeostasis, using the cellular transcriptome as a readout. A bulk RNA sequencing (RNAseq) analysis was performed on epithelial cells isolated from the ileum and colon of SEPT9-KO mice and their control littermates. Principal component analysis (PCA, Figure 5A) illustrated the clustering of the samples based on gene expression profiles, highlighting the distinct transcriptional landscapes in response to the SEPT9 knockout. Loss of SEPT9 resulted in significant downregulation of 75 genes and upregulation of 59 genes in mouse ileal epithelial cells. Interestingly, 123 genes were upregulated and only 19 genes were downregulated in SEPT9 deficient colonic epithelium (Figure 5B and C). The pathway enrichment analysis revealed upregulation of transcripts related to acute phase and inflammatory response and down-regulation of contractile genes in colonic mucosa of SEPT9-KO mice (Figure 5D). In ileal epithelia, loss of SEPT9 increased genes related to cell-matrix adhesions and downregulated microtubule-related genes (Figure 5E). The observed alterations in contractile, microtubule and matrix adhesion pathways are consistent with known septin functions in regulating other cytoskeletal elements and cell interactions with extracellular matrix^17–19^.

**Fig. 5:**
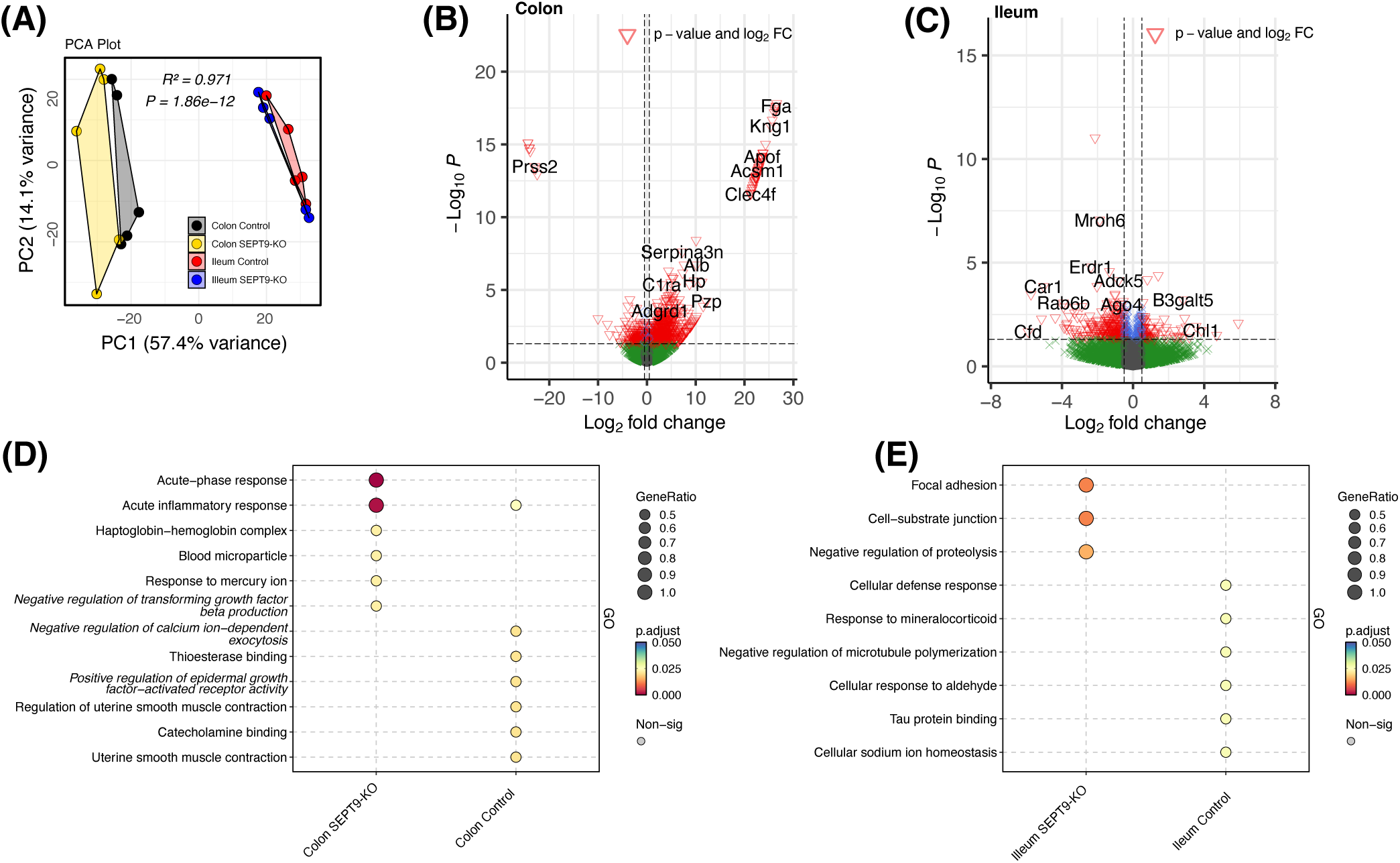
Loss of SEPT9 induces transcription reprogramming of intestinal epithelial cells in mice. (**A**) Principal Component Analysis (PCA) of gene expression profiles in colonic and ileal epithelial cells isolated from SEPT9-KO and control mice. Volcano plots comparing gene expression in (**B**) colonic and (**C**) ileal epithelium of SEPT9-KO and control mice. Significant differentially (p-value and log2FC cut-off) expressed genes are highlighted. Bubble plots representing pathway enrichment analysis of genes with statistically significant differences in expression between the SEPT9-KO and control mice in the (**D**) colonic and (**E**) ileal epithelium. Each bubble corresponds to a specific pathway, with size indicating the gene ratio and color representing the significance of the enrichment. The analysis was performed using isolated IEC from 5 mice/group.

### IEC-specific knockout of SEPT9 exacerbates the severity of acute mucosal inflammation and cell death during experimental colitis

Increased permeability of the intestinal epithelial barrier is known to exacerbate inflammatory response in the gut^4,5,42^. Therefore, we next investigated whether gut barrier disruption observed in SEPT9-KO mice could modulate mucosal inflammation and injury using a dextran sodium sulfate (DSS) model of acute colitis. Exposure to DSS resulted in more severe intestinal disease in SEPT9-KO mice, as compared to their control littermates (Figure 6A and B). The exaggerated disease was characterized by more pronounced body weight loss (Figure 6A) and a significantly higher disease activity index (Figure 6B), which is a composite of body weight loss, occurrence of diarrhea, and intestinal bleeding. Furthermore, SEPT9-KO mice demonstrated higher intestinal permeability to low and high molecular weight tracers on day 7 of DSS colitis (Figure 6C and D). The increased permeability to large (70 kDa) molecules could reflect higher mucosal erosion and IEC loss in SEPT9-KO mice. Surprisingly, a cumulative tissue injury index composed based on the evaluation of H&E-stained sections of distal colon was not significantly elevated in DSS-treated SEPT9-KO mice after 7 days of acute colitis (Supplementary Figure 6). This may reflect leveling of some disease symptoms in SEPT9-KO and control mice at late stages of DSS colitis (Figure 6B).

**Fig. 6:**
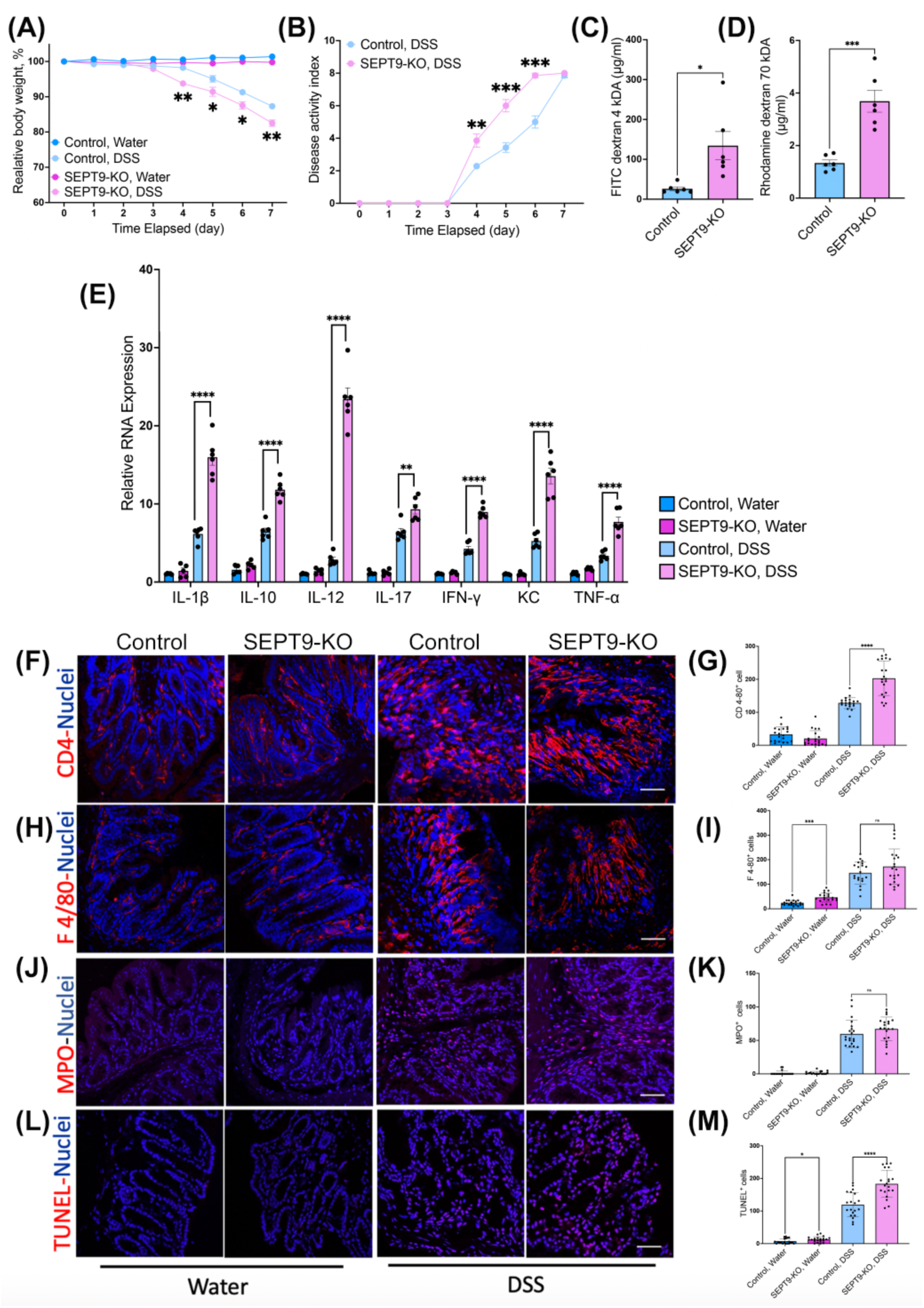
Loss of SEPT9 exacerbates disease severity and inflammatory response during DSS-induced colitis. SEPT9-KO and control mice were exposed to 3% DSS in drinking water, or regular water for 7 days. (**A**) Body weight loss and (**B**) disease activity index, were recorded daily. (**C and D**) Intestinal permeability of DSS-treated SEPT9-KO and control mice. Gut-to-blood passage of 4 kDa FITC-dextran (C) and 70 kDa Rhodamine-dextran (D). (**E**) mRNA expression of inflammatory markers measured in colonic tissues of DSS-treated SEPT9-KO and control mice. **(F-K)** Immunolabeling and quantification of leukocyte infiltration (red) in the colonic mucosa of SEPT9-KO and control mice on day 7 of DSS or water exposure, including T-cells (F and G), monocytes/macrophages (H and I) and neutrophils (J and K), as well as TUNEL labeling of apoptotic cells (L and M). Nuclei are labeled with DAPI (blue). Scale bars= 20 μm. Mean ± SEM (n = 6); *p < 0.05, **p<0.01 *** p<0.001, ****p<0.0001. The scatter dots within the bars in **C, D**, and **E** represent individual mice. The scatter dots in the **G, I, K** and **M** graphs represent pooled cell numbers.

Examining H&E-stained colonic tissue sections provides only a general snapshot of the intestinal architecture. Therefore, we next used a set of specific experimental approaches to evaluate important aspects of mucosal inflammation and tissue injury in DSS-treated animals. Expression of proinflammatory cytokines and chemokines was examined by the quantitative RT-PCR analysis in tissue samples of the distal colon collected on day 7 of DSS administration. mRNA expression of several major cytokines (TNF, IFNψ, IL-1β, IL-10, IL-12, IL-17) and one chemokine, KC, was significantly higher in tissue samples of DSS-treated SEPT9-KO mice, as compared to their control littermates (Figure 6E). Of note, mRNA levels of these mediators were not elevated in colonic tissue of control SEPT9-KO mice without DSS treatment (Figure 6E). This suggest that the increased permeability of the healthy gut barrier is insufficient to trigger spontaneous mucosal inflammation.

Since recruitment and activation of immune cells serve as a major driver of mucosal inflammation, we next examined whether loss of SEPT9 affects leukocyte infiltration into inflamed colonic tissue during DSS colitis. Immunofluorescence labeling of CD4, F4/80, and myeloperoxidase (MPO) antigens was utilized to detect T lymphocytes, monocytes/macrophages, and neutrophils, respectively (Figure 6F–K). The number of monocytes/macrophages, but not T cells or neutrophils, was significantly increased in the normal colonic mucosa of SEPT9-KO mice. While DSS colitis induced marked accumulation of all types in immune cells in inflamed colonic mucosa, only T cell numbers were significantly increased in SEPT9-KO mice compared to controls under DSS treatment (Figure 6F-K). Immune cell infiltration and production of proinflammatory cytokines are known to cause excessive IEC death thereby exaggerating gut barrier disruption and mucosal inflammation^35,43^. A terminal deoxynucleotidyl transferase dUTP nick end labeling (TUNEL) assay was used to visualize dead cells since this technique allows to detect both apoptotic and non-apoptotic types of cell death^44^. Expectedly, DSS administration caused a marked increase in TUNEL-positive cells, indicating increased cell death. The magnitude of this mucosal cell death was significantly higher in DSS-treated SEPT9-KO mice compared to their controls (Figure 6L and M). Together, our findings suggest that loss of SEPT9 in the intestinal epithelium markedly increases sensitivity to experimental colitis by propagating the inflammatory response and cell death in the intestinal mucosa.

### SEPT9 expression and localization are altered in the intestinal mucosa of inflammatory bowel disease patients

Given our findings of increased intestinal epithelial barrier permeability and exacerbated mucosal inflammation in SEPT9-KO mice, we next sought to examine the expression and localization of SEPT9 in the inflamed intestinal mucosa of inflammatory bowel disease (IBD) patients. We performed dual immunofluorescence labeling of SEPT9 and E-cadherin in surgically resected colonic and terminal ileum sections of Crohn’s Disease (CD) patients and colonic sections of ulcerative colitis (UC) patients. Resected colonic sections of patients with diverticulitis and healthy margins of colorectal cancers were used as controls. SEPT9 was found to be enriched at E-cadherin-based junctions in normal intestinal mucosa (Figure 7A, arrows). By contrast, SEPT9 was translocated from IEC junctions into the cytoplasm in CD but not in UC tissues (Figure 7A and B, arrowheads). Furthermore, the general intensity of SEPT9 labeling was decreased in the intestinal epithelium of both CD and UC patients (Figure 7C). We validated these findings using paraffin-embedded colonic biopsies obtained from a different cohort of IBD patients and non-IBD controls. Immunohistochemistry revealed a significant decrease in SEPT9 expression in colonic mucosa from CD and UC patients (Figure 7D and E). Together our findings revealed decreased expression and mislocalization of SEPT9 in the intestinal epithelium of IBD patients.

**Fig. 7:**
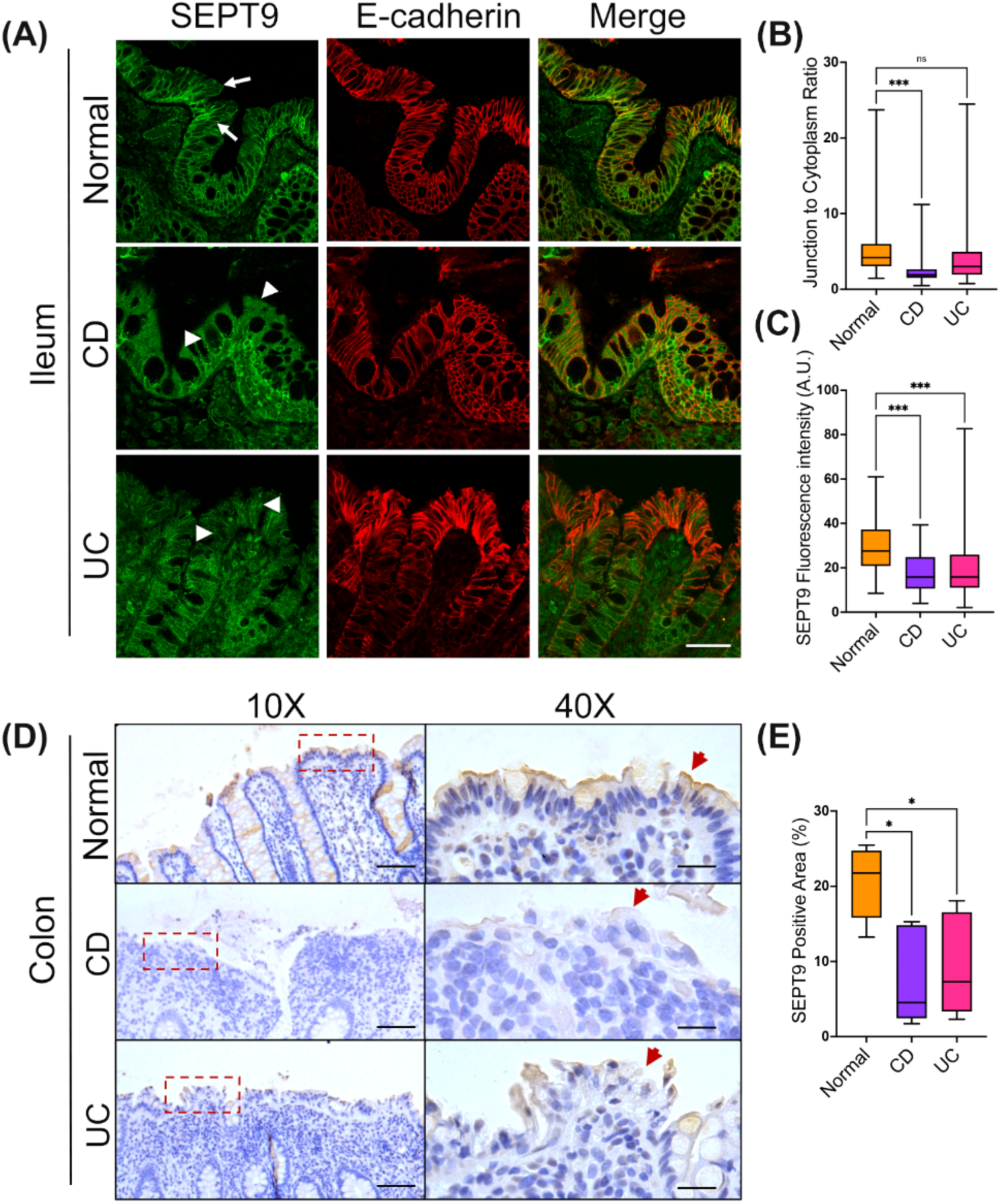
SEPT9 is displaced from junctions and is downregulated in the intestinal epithelium of inflammatory bowel diseases patients. **(A)** Immunofluorescence labeling of SEPT9 (green) and E-cadherin (red) in cryosections of ileal and colonic tissue from IBD patients and non-IBD controls. Arrows point to SEPT9 localization at epithelial junctions in normal sample. Arrowheads indicate mislocalized and decreased SEPT9 labeling in CD and UC tissue samples. Scale bar= 50 μm (**B and C**) Quantification of the junction to cytoplasmic ratio of SEPT9 signal (B) and total SEPT9 signal intensity (C). Mean ± SEM (n= 6 for normal controls and CD patients, and 5 for UC patients). **(D)** Immunohistochemistry of SEPT9 labeling in paraffin sections of colonic tissue sections from control and IBD patients. **(E)** Quantification of total SEPT9 signal intensity. Mean ± SEM (n= 9 for normal controls, 5 for CD and 6 for UC patients). *p<0.05, **p<0.01, ***p<0.001

## DISCUSSION

Increased permeability of the intestinal epithelial barrier is known to exacerbate mucosal inflammation in chronic gastrointestinal disorders, most notably, in IBD^1–3^. IEC barrier integrity depends on proper assembly of junction-associated cytoskeleton, which is composed of elaborate filamentous networks. In this study we identified a novel cytoskeletal mechanism regulating permeability of the gut barrier under homeostatic conditions and during mucosal inflammation. This mechanism involves the SEPT9-based cytoskeleton that strengthens the IEC barrier through the recruitment or scaffolding of NMIIC within the apical junctional actomyosin belt, thereby promoting the assembly and stability of TJs and AJs.

SEPT9 is a unique component of the septin oligomers that self-assemble in high-order cytoskeletal structures with various cellular functions^22,45,46^. Previous studies observed the association of different septins with AJs and reported conflicted effects of septin depletion on AJ protein expression. While SEPT2 upregulated VE-cadherin level in endothelial cells^26^, SEPT2 and SEPT7 knockdown did not affect E-cadherin and β-catenin expression in cultured Caco-2 cells^25^, and SEPT9 depletion decreased E-cadherin and β-catenin levels in MDCK kidney epithelial cells^24^. We show for the first time that SEPT9 associates with TJs in cultured human IEC and mouse intestinal epithelium and, importantly, that loss of SEPT9 increased intestinal epithelial permeability and impaired TJ assembly and dynamics *in vitro* and *in vivo*. In our study, loss of SEPT9 did not affect expression levels of major TJ and AJ protein in human IECs and mouse colonic epithelium (Supplementary Figure 3C). This suggests a scaffolding function of SEPT9 in the intestinal epithelium that involves localized recruitment and/or stabilizing proteins at epithelial junctions.

While SEPT9 may anchor proteins to cell membranes through its phospholipid-binding capability^18^, several lines of evidence suggest that it regulates TJ assembly and function by organizing the peri-junctional actomyosin cytoskeleton. First, our interactome and imaging analyses show that SEPT9 associates with NMIIC motor protein. This interaction is consistent with findings in human breast cancer cells^40^. Secondly, SEPT9 loss resulted in reduced enrichment of junctional NMIIC in IECs, *in vivo* and *in vitro*, coinciding with TJ instability. Finally, NMIIC loss mirrored the barrier-disruptive effects observed with SEPT9 deletion in IECs. NMIIC is highly expressed in differentiated intestinal epithelial cells^35,41^ and is specifically enriched at the peri-junctional actin belt^14,47^. Importantly, NMIIC was previously implicated in regulation of microvilli growth in cultured IECs^41^. However, other epithelial functions of this actin motor remain uncharacterized. Our study provides the first evidence that NMIIC is essential for IEC barrier integrity and is recruited to epithelial TJs through interactions with SEPT9.

Additionally, we observed that SEPT9 loss altered the transcriptional programming of IECs in mice, leading to upregulation of inflammatory pathways in colonic epithelial cells and increase in cell-matrix adhesion pathways in ileal epithelial cells (Figure 5D and 5E). This could be an indirect consequence of enhanced barrier permeability and weakened cell-cell adhesions, which may expose epithelial cells to luminal pathogens and trigger pro-inflammatory signaling, or compensatory upregulation of adhesion-related genes. Alternatively, SEPT9 may directly regulate gene expression, as it was reported recently for the SEPT9-mediated myogenic differentiation of immature myoblasts^48^.

The increased gut permeability in SEPT9-KO animals was not sufficient to cause spontaneous intestinal inflammatory disorders. These data are consistent with previous studies with JAM-A knockout mice^49^ and mice with conditional knockout of β-cytoplasmic actin in IEC^50^ and suggest development of compensatory mechanisms restricting activation of the mucosal immune response under conditions of mildly leaky gut. We found, however, that loss of SEPT9 increases severity of the disease symptoms and enhances mucosal injury and inflammation in acute DSS colitis (Figure 6). While the only previous study reported anti-inflammatory features of septins during Shigella infection in zebrafish^51^, our findings highlight the novel role of the septin cytoskeleton in preventing mucosal damage, gut barrier disruption and inflammation during human IBD-relevant colitis model. These data are in line with previous studies showing exaggerated animal responses to acute colitis in animals deficient in key molecular constituents of the IEC actomyosin cytoskeleton^35,50^. Importantly, our data suggest the dysfunction of the junction-associated septin cytoskeleton in the intestinal epithelium of IBD patients. This is particularly evident in CD, where SEPT9 is downregulated and displaced from epithelial junctions (Figure 7). It is likely that such dysfunction of the junction-association cytoskeleton likely contributes to well-documented gut barrier disruption in IBD patients and could increase severity of the disease.

In conclusion, this study reveals a novel mechanism that regulates integrity of the IEC barrier under homeostatic conditions and plays protective role in the intestinal mucosa during inflammation. The mechanism involves the SEPT9-based cytoskeleton that associates with apical junctions and regulates TJ and AJ assembly and their coupling with the cortical actomyosin belt. Disruption of this junction-associated SEPT9 cytoskeleton, as observed in IBD patients, likely contributes to gut barrier dysfunction and exacerbated mucosal inflammation, hallmarks of these diseases. These findings highlight the critical role of SEPT9 in maintaining intestinal barrier integrity and suggest that targeting the septin cytoskeleton may offer new therapeutic strategies for inflammatory bowel disease.

## Acknowledgments

The authors thank Dr. Ajay Zalavadia and Apryl Helmick from the Lerner Research Institute Digital Imaging and Microscopy Core. Microscopy performed at the Lerner Research Institute Digital Imaging Microscopy Core utilized the Leica SP8 confocal microscope that was purchased with funding from the National Institutes of Health SIG grant 1S10OD019972-01. The authors thank Dr. Min Hui Lim, a Program manager for the Lerner Research Institute Genomic Core for facilitating RNA sequencing. The authors thank Dr. Ernst-Martin Füchtbauer, from University of Aarhus, Denmark for providing SEPT9 floxed mice. The authors thank Dr. Roberto Weigert NCI/NIH for sharing the SEPT9^mNG^. The human samples were provided by Biorepository and Tissue Research Facility at the University of Virginia, and by the Human Tissue Procurement Facility of the Cleveland Clinic through the services of the Biorepository Core funded by National Institutes of Health grant P30 DK097948.

Nayden G. Naydenov and Gaizun Hu contributed equally to this work. Seham Ebrahim and Andrei I. Ivanov are co-senior authors for this study.

## Author Contributions

**S.E.**: Conceptualization, Methodology, Writing – Original Draft, Writing – Review & Editing, Supervision, Project administration, Funding acquisition; **A.I.I.**: Conceptualization, Methodology, Writing – Original Draft, Writing – Review & Editing, Supervision, Project administration, Funding acquisition; **N.G.N.**: Methodology, Validation, Formal analysis, Investigation, Data Curation, Writing – Review and Editing, Visualization; **G.H.**: Methodology, Software, Validation, Formal analysis, Investigation, Resources, Data Curation, Writing – Original Draft, Writing – Review and Editing, Visualization,; **A.Z.**: Formal Analysis, Investigation, Visualization, Writing-Review and Editing; **D.R.**: Methodology, Software, Formal analysis, Resources, Data Curation, Visualization, Writing – Original Draft; **K.M.**: Investigation, Visualization;; **S.L**. Formal Analysis, Investigation, Visualization; **Y.O.**: Investigation; **N.S**.: Formal analysis, Software, Validation, Data Curation, Visualization, Writing-Review and Editing; **G.S.**: Methodology, Software, Formal analysis, Supervision, Writing-Review and Editing; **S.B.**: Formal Analysis, Investigation, Visualization, Writing -Review and Editing; **E.J.**: Formal Analysis, Investigation, Visualization, Writing -Review and Editing; **R.M.**: Software, Formal analysis, Data Curation, Visualization, Writing-Review and Editing; **L.S.**: Formal Analysis, Investigation Writing-Review and Editing; **A.M.M.**: Formal Analysis, Investigation, Writing-Review and Editing; **F.R.**: Methodology, Formal Analysis, Data Curation, Funding acquisition, Writing-Review and Editing.

---

**Fig. S1.**
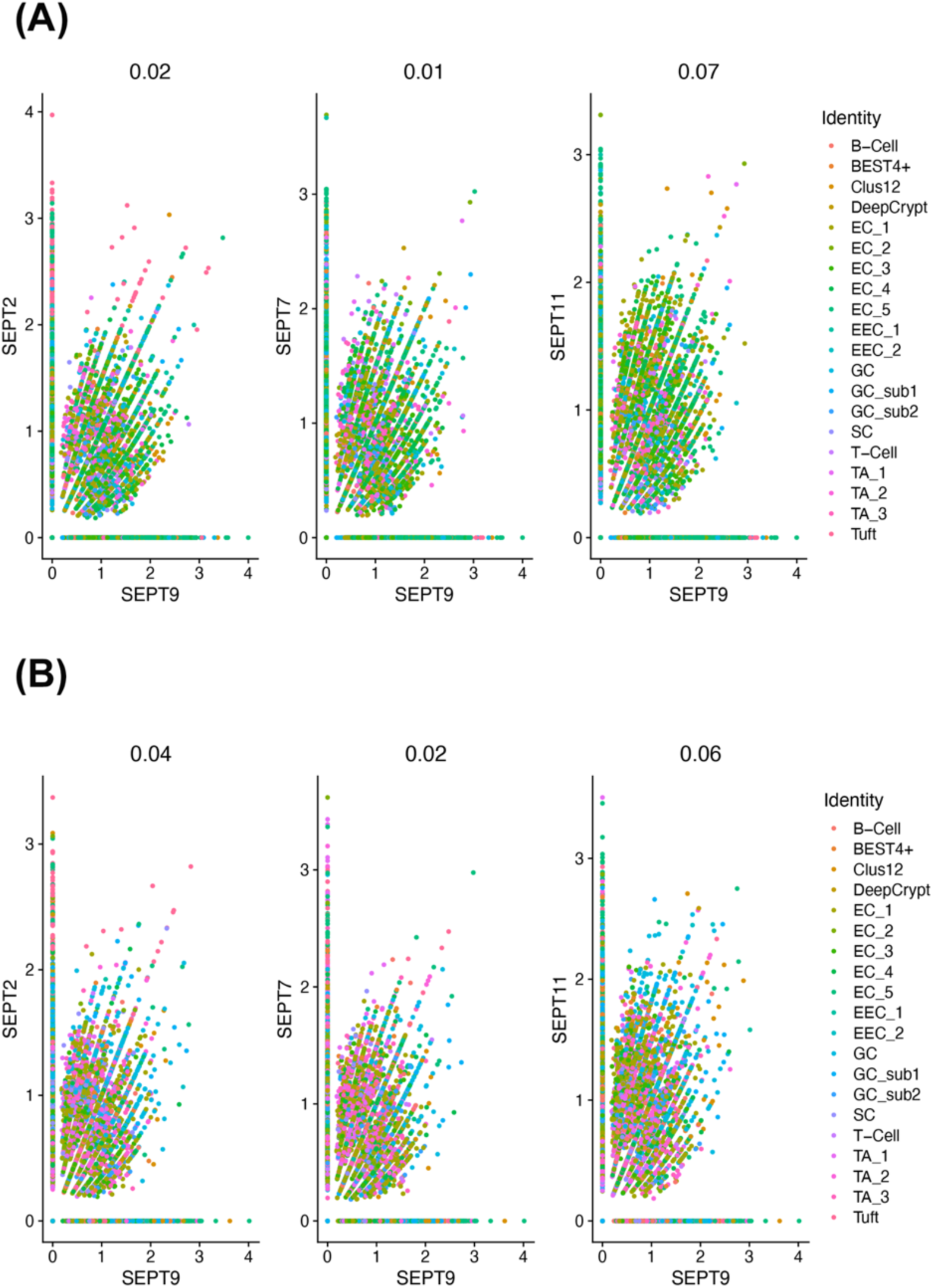
Correlation of SEPT9 expression with expression of other major septin paralogs. The plots show correlation of SEPT9 transcript with the levels of SEPT2, SEPT7 and SEPT11transcripts in epithelial cell subpopulation in normal human ileum **(A)** and normal human colon **(B)**. Numbers of the top of the graphs are Pearson correlation between two genes. Cell cluster abbreviations are: DeepCrypt, deep crypt secretor; EC, enterocytes; EEC, enteroendocrine; GC, Goblet; SC, stem; TA, transit amplifying; Tuft, tuft cells.

**Fig. S2:**
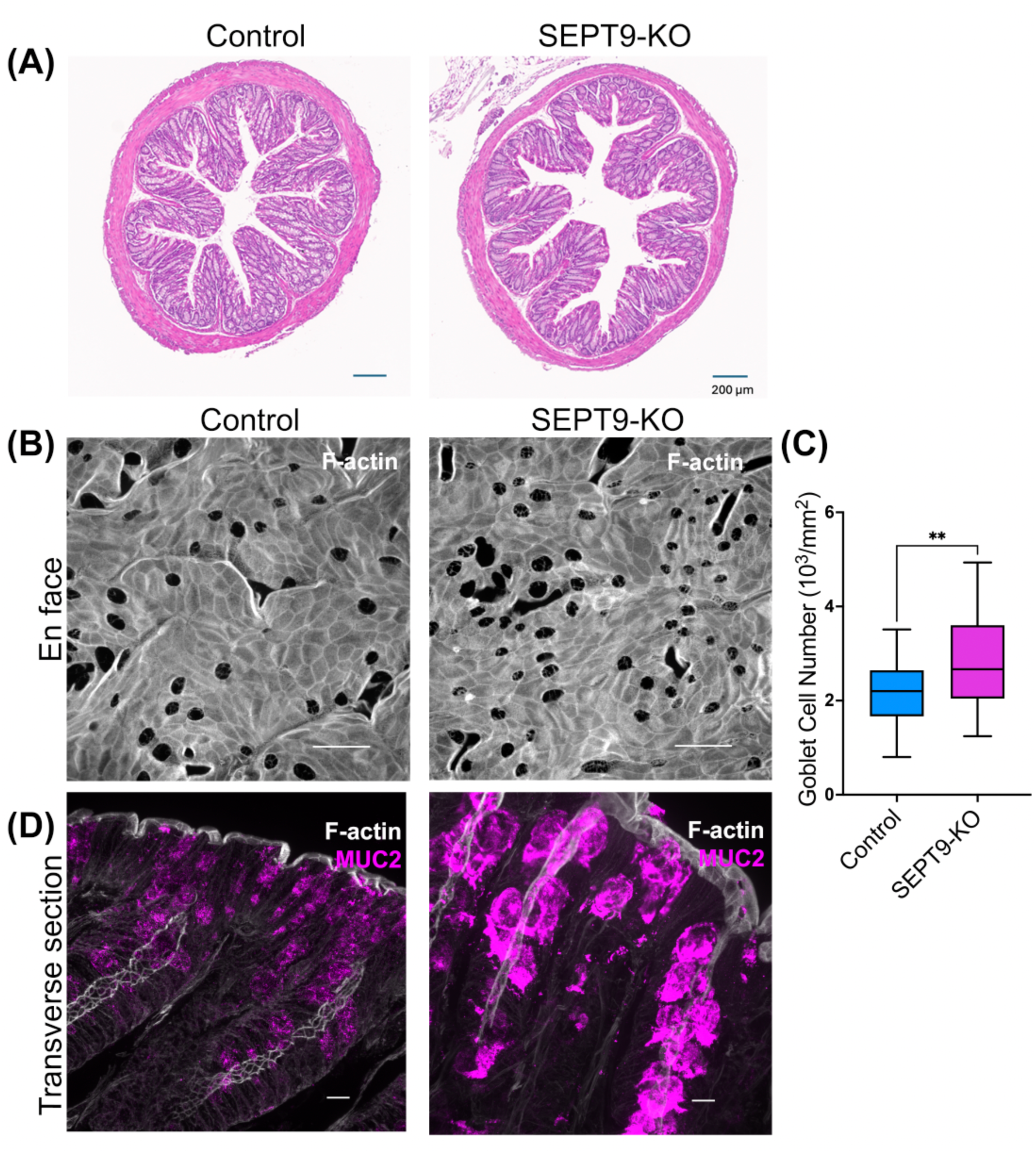
Overall morphology and Goblet cell abundancy in colonic sections on unchallenged control and SEPT9-KO mice. **(A)** Full thickness sections of the distal colonic segments stained with H&E. **(B and C)** *En face* imaging of colonic epithelium labeled with F-actin probe, Alexa Fluor-tagged phalloidin (6 area/mouse, n= 6 mice/group). Representative images (B) and quantification of Goblet cell numbers (dark holes in the F-actin labeling) are shown (C). Data is shown as box whisker plots with whiskers extending to the minimum and the maximum with mean value at the middle of the box body. Student’s t-test. **p<0.01. Scale bar = 25μm **(D)** Dual fluorescence labeling of MUC2 and F-actin in colonic tissues of control and SEPT9-KO mice Scale bar = 10μm.

**Fig. S3:**
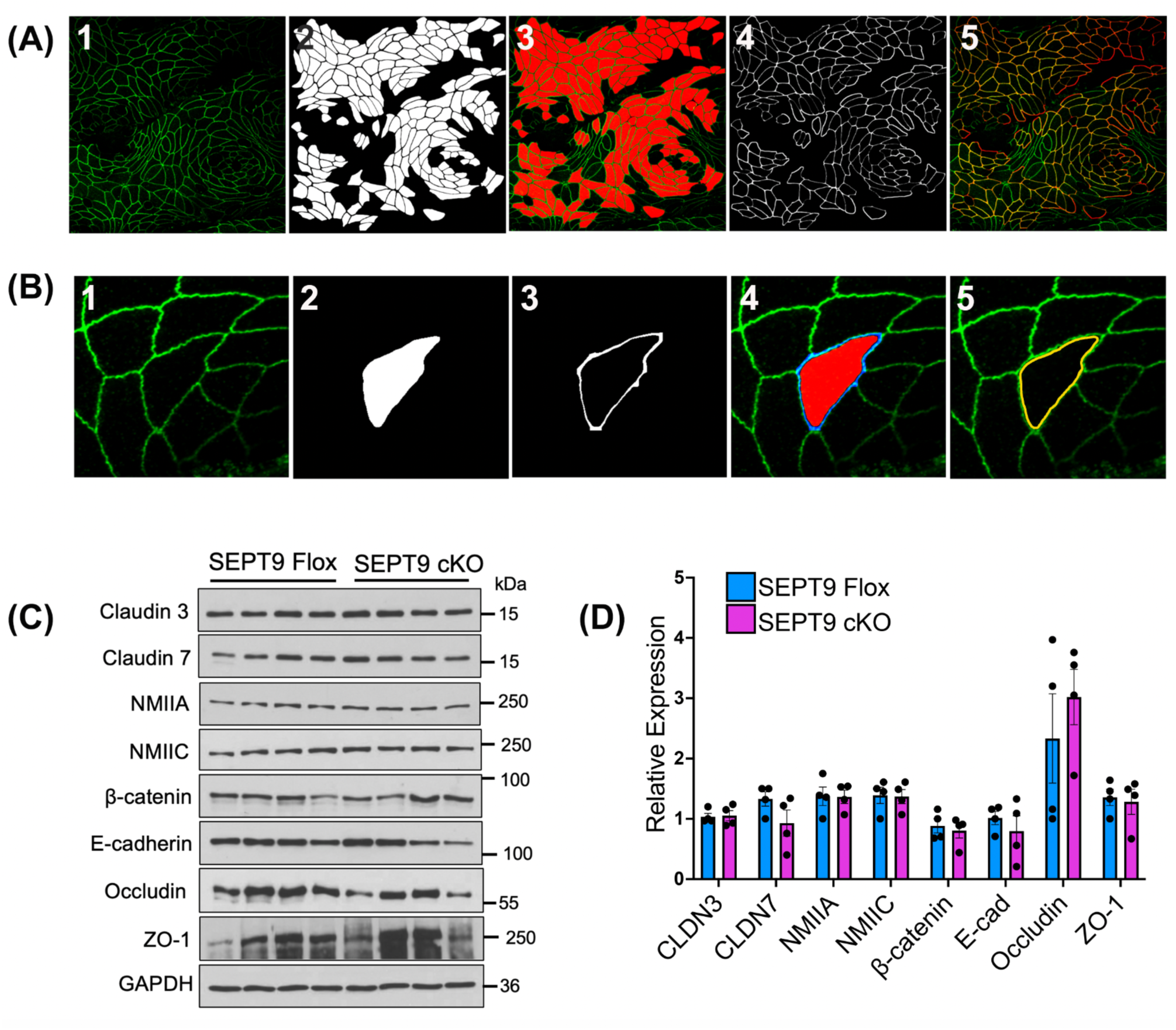
Examples of machine-learning segmentation of colonic epithelium and junctional protein expression in control and SEPT9-KO mice. (A and. **B**) Representative images of *en face* intestinal epithelial segmentation in entire and detailed images. (A1) Representative image of junctional protein quantification. 0.6µm of wholemount colon tissue labeled with ZO-1 antibody was employed as an example (A2) Binary mask of cells generated from the original image with additional manual filtration. (A3) Overlay of the original image (green) with the binary mask (red). (A4) Binary mask of the junctions highlighting the intercellular borders. (A5) Overlay of the original image (green) with the junctional mask (red) in yellow. (B1) Representative image of junctional protein quantification. 0.6µm of wholemount colon tissue labeled with ZO-1 antibody was employed as an example (B2) Binary mask of the cell created from the original image. (B3) Binary mask of the junction of the cell of interest. (B4) Overlay of the original image (green) with the binary mask of the cell (red) and the binary mask of the junction (blue). (B5) Marked area (yellow ring) where junctional FI was measured using ImageJ. (**C and D**) Immunoblotting analysis of junctional protein expression in in colonic scrapes obtained from control SEPT9 Flox and SEPT9 cKO mice. Representative immunoblots (C) and densitometric quantification of protein expression (D) are shown (n= 4/group). The intensities of the bands for each sample were normalized to those for GAPDH, and the intensity of the bands in the controlled group was assigned a value of 1. Data are shown as mean ± SEM. * p<0.05.

**Fig. S4:**
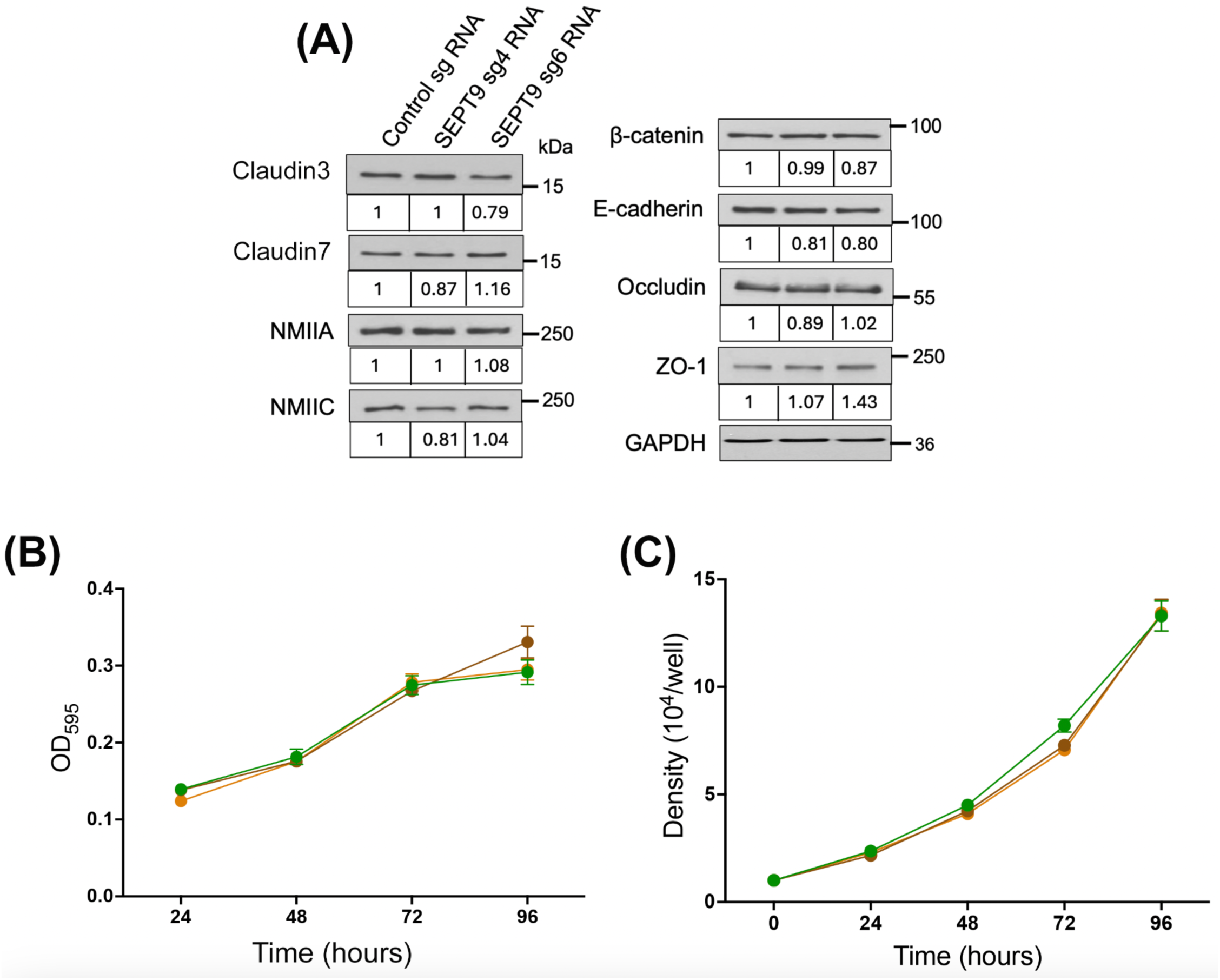
Expression of different junctional and cytoskeletal proteins and proliferation assays of control and SEPT9 KO IEC. **(A)** Immunoblotting analysis of junctional protein and myosin motor expression in control and SEPT9-KO HT-29 cells. Levels normalized to relative expression of GAPDH are shown in the boxes. **(B)** MTT assay and **(C)** cell number counting of control, and SEPT9-KO (sg4: brown; sg6: orange) HT-29 cells at different times after plating. Data shown as mean ± SEM (n =3).

**Fig. S5:**
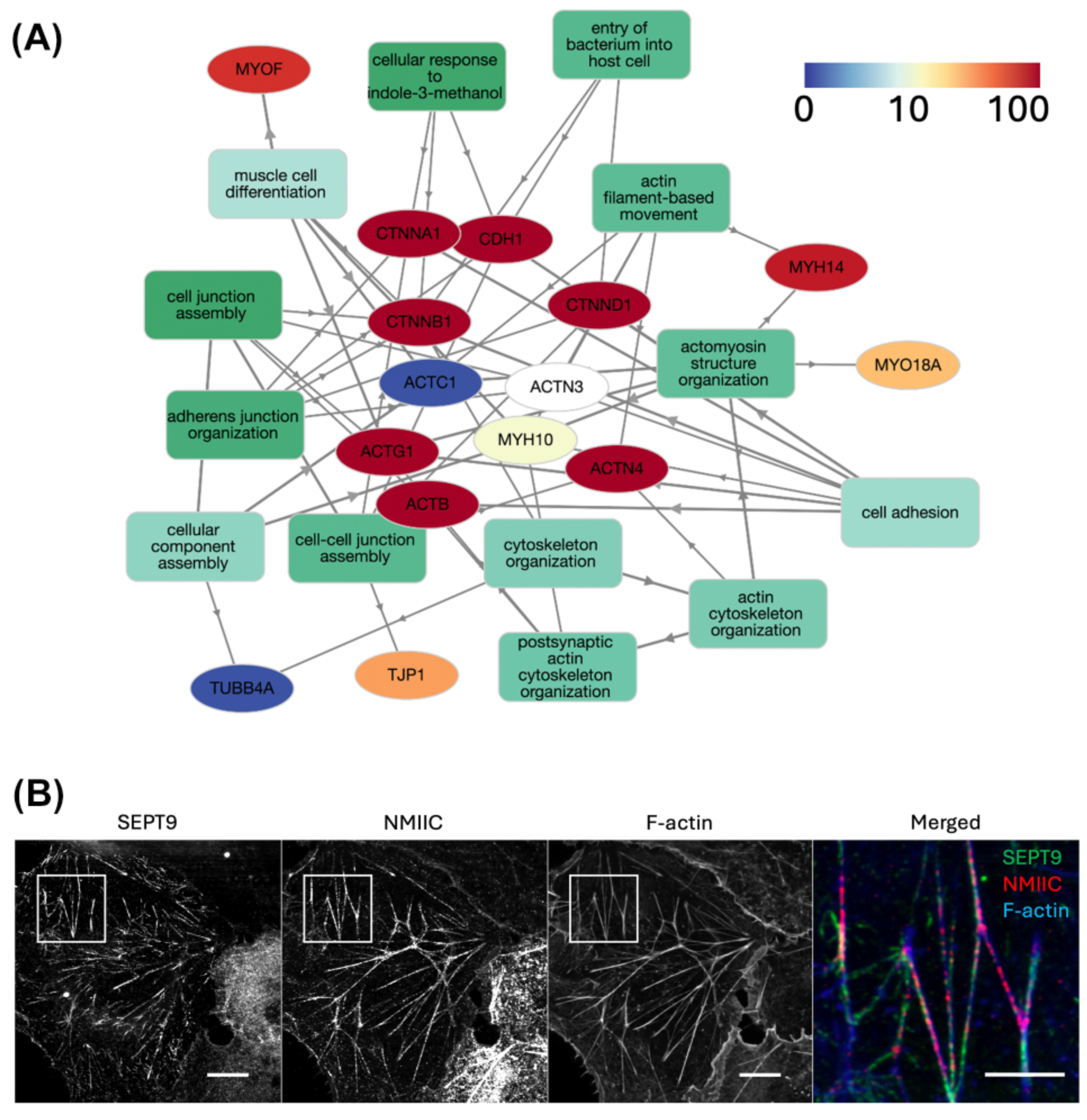
Interactome of SEPT9 and co-assembly of SEPT9 and NMIIC within F-actin bundles. **(A)** Interactome analysis of the binding partners for SEPT9 in IECs *in vitro*. **(B)** Co-transfection of SEPT9 (green) and NMIIC (red) in COS-7 cell line. The white boxes indicated the zoomed area (Merged). Scale bar= 2 μm

**Fig. S6:**
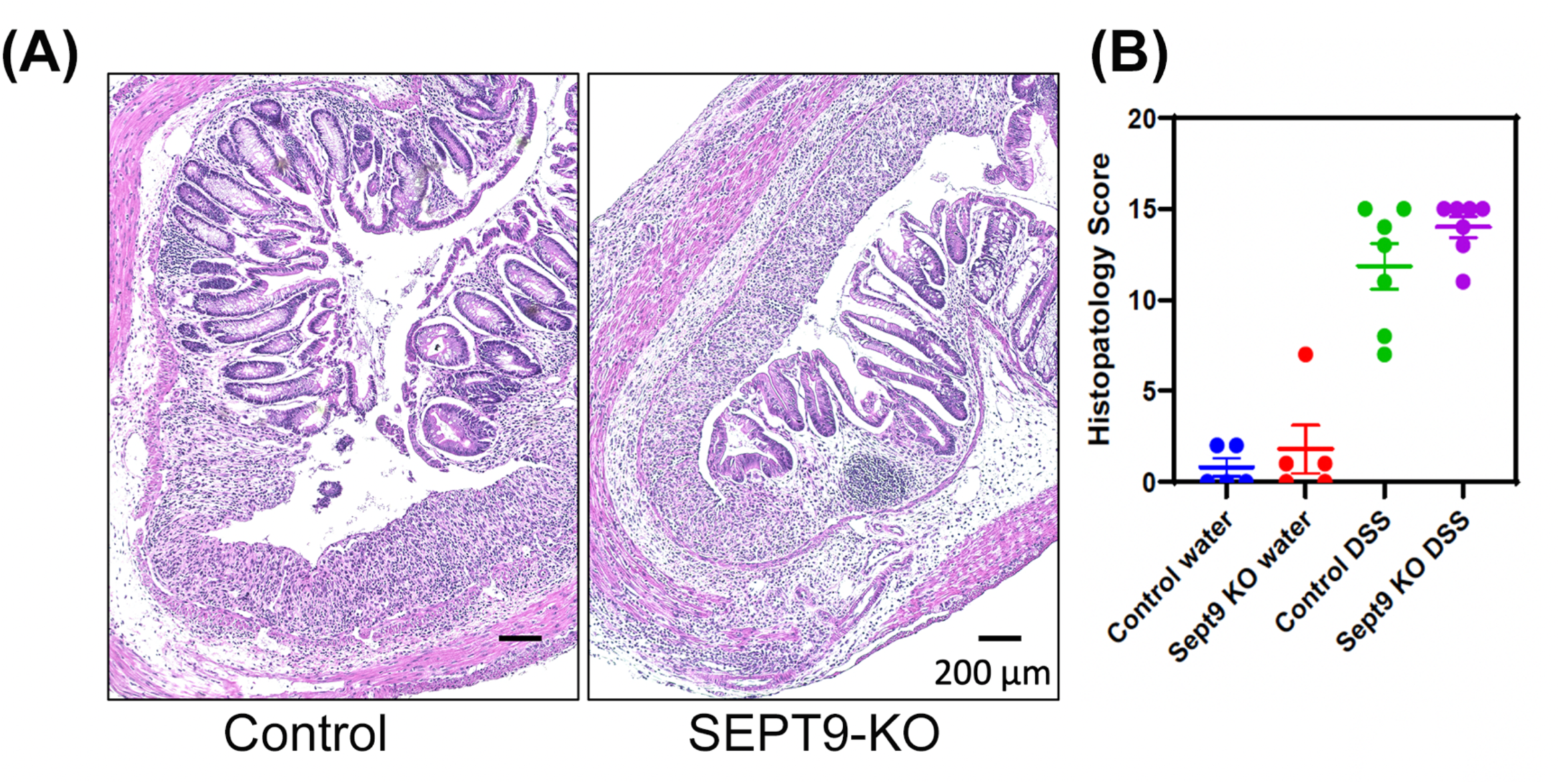
DSS-induced mucosal damage and inflammation in colonic tissue of control and SEPT9-KO mice. Mucosal injury and inflammation were examined in H&E-stained distal colonic sections of Control and SEPT9-KO mice on day 7 of DSS treatment. **(A)** Representative H&E images and **(B)** calculated tissue injury index are shown as mean ± SEM. (n= 5 in water-treated groups and n= 7 in DSS-treated groups)

**Table 1.** Patient information.

| **Cleveland Clinic Patient Cohort** |  |  |  |  |
| --- | --- | --- | --- | --- |
| **Groups** | **Sample number**<br>**(*n*)** | **Gut segment** | **Age, years,**<br>**Mean (range)** | **Male sex, n (%)** |
| Crohn's Disease | 6 | Colon (5) and ileum (1) | 59 (41-69) | 2 (33) |
| Ulcerative colitis | 5 | Colon | 42 (24-57) | 3 (60) |
| Non-IBD control | 6 | Colon | 63 (14-82) | 3 (50) |
| **University of Virginia Patient Cohort** |  |  |  |  |
| **Groups** | **Sample number**<br>**(*n*)** | **Gut segment** | **Age, years,**<br>**Mean (range)** | **Male sex, n (%)** |
| Crohn's Disease | 5 | Colon | 33 (26-45) | 3 (60) |
| Ulcerative colitis | 6 | Colon | 54 (12-83) | 0 (0) |
| Non-IBD control | 9 | Colon | 59 (44-72) | 3 (33) |

## Supplementary Methods

### Antibodies and other reagents

The following primary antibodies were used to detect septins, junctional, cytoskeletal and leukocyte proteins: SEPT9 (Sigma-Aldrich, HPA042564), for immunoblotting and immunofluorescence and (Proteintech, 10769-1-AP,) for immunohistochemistry, ZO-1 (21773-1-AP, Proteintech, for immunoblotting; 33-9100, ThermoFisher, for immunofluorescence), occludin (13409-1-AP, Proteintech), claudin 7 (34-9100, Thermo Fisher Scientific), claudin 3 (34-1700, Thermo Fisher), NMIIA (909801, BioLegend), ZO-1 (40-2200, Thermo Fisher), myeloperoxidase (ab9535, Abcam), mucin 2 (Abcam, ab272692), GAPDH (2118S, Cell Signaling Technology), E-cadherin (AF648, R&D Systems for human tissue immunolabeling, 00101, BiCell for mouse tissue immunofluorescence, PA5-32178, ThermoFisher for human cells immunofluorescence), NMIIC (8189S, Cell Signaling Technology for immunoblotting, PA5-64345, ThermoFisher for immunofluorescence), β-catenin (610153, BD Biosciences) CD4 (550278, BD Biosciences), F4/80 (MCA497GA, Bio-Rad Laboratories) myeloperoxidase (Abcam, Cambridge, UK; ab9535). AlexaFluor-488/555/568/647-conjugated secondary antibodies, were obtained from ThermoFisher Scientific. Horseradish peroxidase-conjugated (HRP) goat anti-rabbit and anti-mouse secondary antibodies were acquired from Bio-Rad Laboratories. Fluorescein isothiocyanate (FITC)-4 kDa dextran (cat# FD4) and Rhodamine B isothiocyanate-70 kDa dextran (cat# R9379) were purchased from Sigma-Aldrich. The ApopTag® Fluorescein in Situ Apoptosis Detection kit was obtained from Sigma-Aldrich (Cat# S7110). All other chemicals were obtained from Sigma-Aldrich or ThermoFisher.

### Generation of SEPT9-mNeonGreen reporter mice

The genetic knock-in mice were generated by inserting the mNeonGreen (mNG) tag into the C-terminus of the SEPT9 sequence. The mice were inbred over three generations, and the fluorescence tag was checked under the fluorescent microscope.

### Generation of Tamoxifen-inducible, Intestinal Epithelium-specific Knockout Mice

Inducible, intestinal epithelium-specific knockout mice (SEPT9^fl/fl^Vil1^CreERT2^, SEPT9-KO) were generated by crossing with SEPT9^fl/fl^ strain (provided by Dr. Ernst-Martin Füchtbauer, University of Aarhus, Denmark) and Vil1^CreERT2^ strain (The Jackson Laboratory, Bar Harbor, ME, stock # 020282). For the SEPT9^fl/fl^ mice, a loxP site was inserted upstream of exon 2. An FRT-flanked neomycin resistance cassette with a 5’ loxP site was inserted downstream of exon 5. Flp-mediated recombination removed the neomycin resistance cassette and left exons 2 through 5 floxed^1^. Vil1^CreERT2^ transgenic mice express a tamoxifen-inducible Cre expression driven by the 9-kb mouse Vil1 promoter. When crossed with a strain containing a LoxP site flanked sequence of interest, the offspring are useful for generating tamoxifen-induced, Cre-mediated targeted deletions. Tamoxifen induces Cre recombination in the intestinal epithelia of the duodenum, jejunum, ileum, proximal and distal colon^2^. Tamoxifen administration was carried out according to the established protocol^3^, and the efficiency of SEPT9 knockout was validated by immunoblotting of colonic epithelial scrapes obtained from SEPT9-KO and control animals. The animal colonies were maintained under pathogen-free conditions in the vivarium of the Lerner Research Institute of Cleveland Clinic and University of Virginia School of Medicine. Standard feed and tap water were available, *ad libitum*. The mouse room was on a 12 h light/dark cycle (lights on at 7:00 A.M.). At the beginning of colitis experiments, mice weighed 18-25 g, with no meaningful difference between the body masses of mice of different genotypes. All procedures were conducted under an animal research protocol approved by the Lerner Research Institute (IACUC protocol # 00001872) and University of Virginia Animal Care and Use Committees (Protocol# 4382) in accordance with the National Institutes of Health Animal Care and Use Guidelines.

### Induction and characterization of dextran sodium sulfate (DSS) colitis

Experimental colitis was induced in 8–10-week-old SEPT9-KO mice and control SEPT9 flox littermates by administering a 3% (w/v) solution of DSS (molecular weight range 35,000-50,000 kDa; Thermo Fisher, Cat# J14489.22) in drinking water. Unchallenged animals received tap water *ad libitum*. Both male and female mice were used at roughly equal numbers. Animals were weighed daily and monitored for signs of intestinal inflammation. The disease activity index was calculated as previously described, by averaging numerical scores of body weight loss, stool consistency, and intestinal bleeding^4^. With regards to body weight, no weight loss was scored as 0, loss of 1-5% was scored as 1; 5-10% weight loss as 2; 10-15% as 3, and more than 15% weight loss was scored as 4. For stool consistency, well-formed pellet was scored as 0, soft and semi-formed stool as 2, and liquid stool or diarrhea scored as 4. For intestinal bleeding, no blood was scored as 0, hemoccult-positive stool as 2, and gross rectal bleeding was scored as 4. On day 7 of DSS administration, animals were euthanized, with their colonic tissue harvested and separated into several segments. The samples were either fixed in a 4% formaldehyde solution, snap-frozen in liquid nitrogen or embedded in an optimal cutting temperature medium and snap-frozen for subsequent histological and biochemical examination. Formalin-fixed samples were paraffin-embedded, sectioned, and stained with hematoxylin and eosin (H&E). The tissue injury index was calculated based on microscopic examination of H&E sections, as previously described^5^. The index represents the sum of individual scores reflecting submucosal edema, crypt hyperplasia/inflammation, leukocyte infiltration, and epithelial erosion.

### Measuring intestinal permeability in vivo using fluorescently labeled tracers

The in vivo intestinal permeability assay was performed in SEPT9-KO and Control mice following the method described previously^6^. Food-deprived animals were given a mixture of FITC-labeled dextran (4 kDa; 80mg/100 g body weight) and rhodamine-labeled dextran (70 kDa; 40mg/100g body weight) dissolved in phosphate-buffered saline (PBS). Control mice received PBS only. Animals were euthanized 3 hours later for blood collection via cardiac puncture, blood serum was obtained by centrifugation, and FITC and Rhodamine fluorescence intensities were measured using a Biotech Synergy H1 plate reader (Agilent Technology, Santa Clara, CA) with excitation and emission wavelengths at 495/525nm and 555/585nm, respectively. The fluorescence intensity of dextran-free serum was subtracted from each measurement. The concentration of fluorescent tracers in blood serum was calculated using the standard curves prepared via serial dilutions of stock solutions of FITC-dextran and Rhodamine-dextran in PBS.

### Cell culture

HT-29 clone cF8, a well-differentiated clone of HT-29 human colonic epithelial cells^7,8^ were provided by Dr. Judith M Ball (College of Veterinary and Biomedical Sciences, Texas A&M University, College Station, TX). DLD1 (Cat# CCL-221) and Caco-2BBE (Cat# CRL-2102) human colonic epithelial cells were obtained from the American Type Culture Collection (ATCC, Manassas, VA). HT-29, DLD1 and Caco-2BBE cells were cultured in DMEM (ATCC, Cat# 30-2002) medium supplemented with 10% fetal bovine serum (FBS), HEPES, non-essential amino acids, and penicillin-streptomycin antibiotic. The cells were seeded on collagen-coated Transwell filters (Falcon, Corning, NY, Cat# 353095), or glass coverslips for permeability measurements and immunolabeling experiments; and on 6-well plastic plates for other functional and biochemical studies. All cultured cells were mycoplasma-free according to the mycoplasma PCR detection assay (PromoCell, Heidelberg, Germany, Cat# PK-CA20-700-20).

### Cell proliferation assay

The cell proliferation profiles were measured by either cell counting using hemocytometer or by the MTT assay using the MTT cell growth assay kit (Sigma-Aldrich, Cat#CT01) according to the manufacturer instructions. The MTT optical density was measured using the Biotech Synergy H1 plate reader. The cells numbers and MTT absorbance were determined daily for 4 days after cell plating.

### Quantitative real-time RT-PCR analysis of inflammatory gene expression in mouse colonic tissues

Total RNA was isolated from distal colonic segments of SEPT9-KO and Control animals using a RNeasy mini kit (Qiagen) followed by DNase treatment to remove genomic DNA. Total RNA (1 µg) was reverse transcribed using an iScript cDNA synthesis kit (Bio-Rad Laboratories). Quantitative real-time RT-PCR was performed using iTaq Universal SYBR Green Supermix (Bio-Rad Laboratories) in a CFX96 Real-time PCR System (BioRad). The threshold cycle number (Ct) for specific genes of interest and a housekeeping gene was determined based on the amplification curve representing a plot of the fluorescent signal intensity versus the cycle number. The relative expression of each gene was calculated by a comparative Ct method based on the inverse proportionality between Ct and the initial template concentration (2-ΔΔCt), as previously described^10^. This method is based on two-step calculations of ΔCt = Cttarget gene - CtGAPDH and ΔΔCt = ΔCte-ΔCtc, where index e refers to the sample from any DSS or water-treated SEPT9-KO, or Control mice, and index c refers to the sample from a water-treated control animal assigned as an internal control. The primer sequences are described in our published study^11^

### Immunoblotting analysis

Mouse colonic segments were harvested, longitudinally dissected, opened, and washed with ice-cold PBS. Epithelial cells were collected by gently scraping the exposed interior with razor blades, then snap frozen in liquid nitrogen for further analysis. Intestinal epithelial scrapings were lysed and homogenized in RIPA buffer containing a protease inhibitor cocktail, phosphatase inhibitor cocktails 2 and 3, and phenylmethylsulfonyl fluoride (all from Sigma-Aldrich). Samples were diluted with the 2x SDS sample loading buffer and boiled. Total lysates of cultured human IEC cells were prepared similarly to the mouse tissue lysates. SDS-polyacrylamide gel electrophoresis was conducted using standard protocols with an equal amount of total protein loaded per lane (10 µg), followed by transfer to a nitrocellulose membrane at 4°C for either 1 or 12 h depending on protein size. After transfer, the membranes were blocked in 5% TBST-20 buffered skim milk solution for 1 h at room temperature and then incubated with primary antibodies either for 1 hour at room temperature or overnight at 4°C. Primary antibodies were used at 1:500 or 1:1,000 dilution. Membranes were washed, incubated with HRP-conjugated secondary antibodies (1:10,000 dilution) at room temperature, and the labeled proteins were visualized using standard enhanced chemiluminescence reagents (Sigma-Aldrich) and X-ray films. Protein expression was quantified via densitometry using ImageJ 1.51K software (National Institutes of Health, Bethesda MD). Signal intensities of each protein were normalized by the signal intensity of a housekeeping protein, GAPDH, in the same sample. The data are presented as normalized values, where expressions of either a reference control mouse sample or a human control sgRNA sample were taken as 1.

### Affinity mass spectrometry of SEPT9 interactome

DLD-1 cells were grown to sub-confluency and transfected with human SEPT9-GFP plasmid (OriGene, Rockville, MD; RG225682). The transfected cells were harvested 24 h after transfection and lysed. SEPT9-GFP protein was purified by coimmunoprecipitation, digested with trypsin, and desalted using ChromoTek iST GFP-Trap IP-MS sample preparation kit (Proteintech, Cat gtak-iST). Desalted samples were analyzed in duplicate by nanoLC-MS/MS using a Dionex Ultimate 3000 (Thermo Fisher Scientific, Bremen, Germany) coupled to an Orbitrap Eclipse Tribrid mass spectrometer (Thermo Fisher Scientific, Bremen, Germany). Peptides was loaded onto an Acclaim PepMap 100 trap column (300 μm × 5 mm × 5 μm C18) and gradient-eluted from an Acclaim PepMap 100 analytical column (75 μm × 25 cm, 3 μm C18) equilibrated in 96% solvent A (0.1% formic acid in water) and 4% solvent B (80% acetonitrile in 0.1% formic acid). The peptides were eluted into the mass spectrometer at 300 nL/min up to 90% B over a period of 2 hours. MS2 spectra were acquired using a data-dependent instrument method with the following settings: positive ion mode was used with 2.0 kV at the spray source, RF lens at 30% and data dependent MS/MS acquisition with XCalibur version 4.3.73.11. Full MS scans were acquired in the Orbitrap from 375 to 1500 *m*/*z* with 120,000 resolution. Data dependent selection of precursor ions was performed in Cycle Time mode, with three seconds in between Master Scans, using an intensity threshold of 2e4 ion counts and applying dynamic exclusion (*n* = 1 scans within 30 s for an exclusion duration of 60 s and ±10 ppm mass tolerance). Monoisotopic peak determination was applied, and charge states 2–6 were included for HCD MS2 scans (quadrupole isolation mode; 1.6 *m*/*z* isolation window, normalized collision energy at 30%). The resulting fragments were detected in the Orbitrap at 15,000 resolution with standard AGC target and dynamic maximum injection time mode. Raw MS data was searched using Proteome Discoverer against a custom fasta database that contained all Gencode v41 protein sequences as well as SEPT9 isoform sequences. The following parameters were used: trypsin with maximum 2 missed cleavage sites, 10 ppm precursor mass tolerance and 0.02 Da fragment ion mass tolerance, dynamic amino acid modifications (Oxidation / +15.995 Da (M) and Phospho / +79.966 Da (S, T, Y)) and dynamic n-terminal modifications (Acetyl / +42.011 Da, Met-loss / -131.040 Da (M), and Met-loss+Acetyl / -89.030 Da (M)). A static modification was used on Cysteine (Carbamidomethyl / +57.021 Da).

### Examining leukocyte infiltration and cell death in mouse tissues by fluorescence labeling and confocal microscopy

Immunofluorescence labeling of the T-cell marker, CD4, the macrophage marker, F4/80, and the TUNEL assay were performed using frozen sections of mouse colonic mucosa. The sections were fixed with absolute ethanol at -20°C for 20 minutes and rinsed 3 times in cold PBS. The sections were blocked for 60 minutes at room temperature in Hanks HEPES-buffered salt solution containing 1% bovine serum albumin, followed by overnight incubation at 4°C with primary antibodies. Samples were then washed and incubated with Alexa dye-conjugated secondary antibodies for 60 minutes, then rinsed with blocking buffer. The TUNEL labeling of frozen tissue sections was performed using the ApopTag Fluorescein in Situ Apoptosis Detection Kit, according to the manufacturer’s instructions. For MPO staining, formalin-fixed paraffin embedded tissue sections were deparaffinized following by the antigen retrieval using a R-Universal Epitope Recovery Buffer (Electron Microscopy Sciences Hatfield, PA, Cat# 62719-20). The sections were blocked for 60 minutes in 1% bovine serum albumin in Hanks HEPES-buffered and stained as described above. All labeled samples were mounted on slides using ProLong Antifade mounting reagent with DAPI (Thermo Fisher Scientific, Cat #P36941). Fluorescently labeled tissue sections were imaged using Leica HCX PL APO 40xPH3CS (1.25 NA) OIL immersion objective and Leica TCS SP8 AOBS confocal laser scanning system attached to a Leica DMi8 inverted epifluorescence microscope (Wentzler, Germany). The Alexa Fluor 488 and 555 signals were acquired sequentially in frame-interlace mode, to eliminate cross talk between channels.

### Immunofluorescence labeling of SEPT9 in human intestinal tissue samples

Surgically resected full-thickness sections of human intestine were fixed in 10% neutral-buffered formalin for 24h. After fixation, formalin was replaced with a 10% sucrose solution for 2h and dehydration was continued by sequential incubation of the samples with 20% and 30% sucrose solutions for 2h each. Finally, the samples were transferred into 50% OCT/30% sucrose solution, embedded into OCT and snap frozen on dry ice. The frozen blocks were used to prepare 10 µm sections by using a Leica cryostat and the sections were processed for immunofluorescence labeling. Briefly, frozen sections were permeabilized with 0.5% of Triton-X100 for 5 min at room temperature, blocked for 60 min in a blocking buffer (PBS containing 1% bovine serum albumin, pH 7.4) followed by 60 min incubation with anti-SEPT9 rabbit polyclonal antibody and anti E-cadherin goat polyclonal antibody diluted in the blocking buffer (1:200 dilution). Afterwards, the samples were washed three times with the blocking buffer, incubated with Alexa-Fluor-488–conjugated donkey anti-rabbit and AlexaFluor-647–conjugated donkey anti-goat secondary antibodies at 1:1,000 dilution in the blocking buffer for 60 min, rinsed three times with the blocking buffer, and mounted on slides with ProLong™ Gold Antifade mounting medium with DAPI. Immunofluorescence labeled tissues were imaged using Leica 100x CS2 (1.4NA) OIL immersion objective and Leica TCS SP8 AOBS confocal laser scanning system attached to a Leica DMi8 inverted epifluorescence microscope (Wentzler, Germany). The Alexa Fluor 488 and 647 signals were acquired sequentially in frame-interlace mode, to eliminate cross-talk between channels.,

### Immunohistochemical (IHC) labeling of SEPT9 in paraffin embedded samples

Harvested human colon samples were paraffin-embedded, sectioned at 5μm thickness, and stained for SEPT9 (Proteintech, 10769-1-AP,) as described previously^12^. Briefly, the slides were blocked using 10% normal goat serum and incubated with the primary antibodies overnight at 1:200 dilution at 4°C. The slides were incubated with a biotinylated universal secondary antibody and then streptavidin-peroxidase complex (Vector Labs PK-7800,) followed by staining with 3,3′-diaminobenzidine (34002, Thermo Scientific) and counter-staining with hematoxylin (Sigma-Aldrich, 51275,). Finally, the finished slides were dehydrated in gradient ethanol and xylene and mounted in a mounting medium. An optical microscope captured the non-overlapping images from the intestinal mucosa. The percentage of positively stained area for the markers was determined using ImageJ (version 2.0.0, National Institutes of Health).

### Bulk RNA sequencing of mouse intestinal epithelial cells

Ileal and colonic epithelial cells were isolated from SEPT9-KO and control Control mice by the sequential dithiothreitol/EDTA extraction protocol as described previously. Total RNA was extracted using the Qiagen RNeasy Plus Mini kit (Qiagen, Cat#74134), using a modified version of the manufacturer’s protocol. Sample lysates were added to QIAShredder columns and centrifuged, and the homogenized lysates were added to gDNA eliminator columns. The columns were centrifuged, and the lysates were combined with ethanol; the samples were then added to RNeasy Plus Mini spin columns and centrifuged. The membranes were washed six times with RPE buffer and centrifuged after each wash. RNA was eluted using 25 μl of nuclease-free water. The extracted RNA concentrations and the 260/280 absorption ratio were measured using the Qubit assays and Nanodrop. The Total RNA libraries were prepared in the LRI Genomic Core following the manufacturer’s protocol (Illumina, Document #1000000124514). In brief, ribosomal RNA was depleted, and the RNA underwent fragmentation and denaturation. Subsequently, cDNA was synthesized, followed by end repair, adenylation, and adaptor ligation, culminating in PCR amplification. The libraries were evaluated for quality, quantified, and sequenced at 20 million reads per sample on the Illumina sequencer Novaseq 6000.

### Bioinformatic analysis of bulk RNAseq data

Raw sequencing reads were initially assessed for quality using FastQC (FastQC v0.12.1). Following quality assessment, reads were quality-trimmed to remove low-quality bases and adapter sequences using Trimmomatic (version v0.39). The parameters used for trimming were set as follows: a minimum quality score of 20 and a sliding window of 4 bases. The trimmed reads were subsequently re-evaluated for quality using FastQC to confirm that all quality metrics met the acceptable thresholds. To eliminate contaminating host (GRCm39 (Mouse Genome Reference Consortium, Version 39) reads, we utilized BBMap (version 38.99). This step involved aligning the trimmed reads against a mouse reference genome (source) to remove any reads originating from the host. The “filterbyname” option retained only non-host reads for downstream analysis. This ensured that the subsequent mapping and analysis primarily focused on the target transcriptome without interference from host sequences. The cleaned reads were mapped to the mouse genome using the STAR aligner (version 2.7.10.A). STAR was chosen for its speed and accuracy in handling RNA-seq data. Mapping parameters were optimized to include two-pass mapping to improve splice site detection and reduce the number of multi-mapping reads. The output from STAR included aligned reads in BAM format. The count matrix was generated from the STAR output using the featureCounts function from the Subread package in R (version 2.0.1). This step involved using the mouse genome’s corresponding gene annotation file (GTF format). The featureCounts summarized the number of reads mapping to each gene, which was crucial for subsequent differential expression analysis. Differential abundance analysis was conducted using the DESeq2 package in R (version 1.38). The raw count data was normalized for differences in sequencing depth and library composition using the median of ratios method implemented in DESeq2. A negative binomial generalized linear model was used to identify differentially expressed genes (DEGs), and genes with an adjusted p-value of < 0.05 and a fold change ≥ 2 were considered significant. Pathway enrichment analysis was performed using the enrichR R package. Enrichment analysis was conducted with several pathway databases, including, GO, KEGG and Reactome, to identify biologically relevant pathways associated with the identified DEGs. The top enriched pathways were selected based on a p-value threshold of < 0.05. Results were visualized using ggplot2 (version 3.4.2), a powerful R package for data visualization. Key visualizations included heatmaps for normalized count data of the top DEGs, volcano plots depicting the significance and magnitude of expression changes, and pathway analysis plots illustrating the enriched biological processes. The sequence data were uploaded to the NCBI GEO database. Submission ID: SUB14911003; BioProject ID: PRJNA1195935.

### Single cell RNAseq data analysis

Single-cell RNA alignment files from 6 non-IBD patients (paired ascending colon & terminal ileum) were downloaded from GEO database (GSE242087). Samples were aligned to the human reference genome (GRCh38-2020-A) using CellRanger v7.0.1. All downstream analysis was performed using Seurat v5.1.0 in R (v4.3.1). Cells with mitochondrial ratios less than 40% (removes low-quality/dying cells) and total RNA transcript counts less than 50k were removed (removes potential multi-plets). The remaining cells from each sample were normalized for sequencing depth using the NormalizeData function and variable features from each sample were identified using the FindVariableFeatures function. To reduce computational time and memory usage, a ‘sketch’ assay was generated using the top 5,000 candidate cells from each sample by calculating a ‘LeverageScore’ using the SketchData function. The ‘sketch’ assay was then integrated together to remove batch effects using the standard integration pipeline in Seurat to create a UMAP. A total of 19 clusters were identified that included immune cells & fibroblasts (11 clusters, deemed ‘Other’) and standard epithelial lineage cells (8 clusters, deemed ‘Epithelial’ by predominant expression of EPCAM). Marker genes for each cluster were determined using the wilcoxauc function from the ‘presto’ package (v1.0.0). Both the integrated UMAP coordinates and cluster identification calculated for the ‘sketch’ assay were projected onto the full dataset for subsequent analysis. For further refinement and identification of epithelial subtypes, the 8 ‘Epithelial’ clusters were subset off and ran through the same analysis pipeline, which resulted in 20 unique clusters.

### Quantitative Image Analysis

An open-source software QuPath v.0.3.2 ^15^was used to quantify the number of F4/80, CD4, MPO and TUNEL positive cells. Whole microscopic images (at × 40 magnification) were imported to QuPath, and positive cells were detected using optical density sum in the red and DAPI channels of the obtained images. For each type of labeling, the numbers of the red channel positive cells were counted in 5 different images per animal, and tissue sections of 5-7 different animals per each experimental group were examined. The animal numbers for each experimental group are presented in the figure legends. To quantify the SEPT9 signal intensity at the cell-cell junctions, the images were imported into ImageJ 1.51K, and a small rectangular area manually drawn over the junctional contacts labeled by E-cadherin (red). The selected area was transferred over the SEPT9 image (green). The signal intensity of SEPT9 at the cell-cell contact junctions was measured along with signal intensity of the same rectangular area of the adjacent cytoplasmic region. The fluorescence intensity ratio between cell-cell junctions and the cytoplasm was calculated. To calculate the overall intensity of the SEPT9 signal, fluorescence intensities of the paired junctional and cytoplasmic areas were averaged. Ten different measurements were taken per each confocal image, and three different images per each tissue sample were analyzed. A one-way ANOVA with Bonferroni’s post hoc test was used (to compare CD and UC patients; with the Normal controls). p values < 0.05 were considered statistically significant and all statistical analysis was performed using GraphPad Prism 10.

